# LMO2 is critical for early metastatic events in breast cancer

**DOI:** 10.1101/2021.05.26.443198

**Authors:** Shaheen Sikandar, Jane Antony, Gunsagar S. Gulati, Angera H. Kuo, William Hai Dang Ho, Soumyashree Das, Chloé B. Steen, Thiago Almeida Pereira, Dalong Qian, Philip A. Beachy, Fredrick Dirbas, Kristy Red-Horse, Terence H. Rabbitts, Jean Paul Thiery, Aaron M. Newman, Michael F. Clarke

**Author notes:** These authors contributed equally (Shaheen Sikandar, Jane Antony, Gunsagar S. Gulati). Co-senior authors (Aaron M. Newman, Michael F. Clarke).

## Abstract

Metastasis is responsible for the majority of breast cancer-related deaths, however identifying the cellular determinants of metastasis has remained challenging. Here, we identified a minority population of immature *THY1*^+^/*VEGFA*^+^ tumor epithelial cells in human breast tumor biopsies that display angiogenic features and are marked by the expression of the oncogene, *LMO2*. Higher abundance of *LMO2*^+^ basal cells correlated with tumor endothelial content and predicted poor distant recurrence-free survival in patients. Using *MMTV-PyMT/Lmo2*^*CreERT2*^ mice, we demonstrated that *Lmo2* lineage-traced cells have a higher propensity to metastasize. LMO2 knockdown in human breast tumors reduced lung metastasis by impairing intravasation, leading to a reduced frequency of circulating tumor cells. Mechanistically, we find that LMO2 binds to STAT3 and is required for STAT3 activation by TNFα and IL6. Collectively, our study identifies a population of metastasis-initiating cells with angiogenic features and establishes the LMO2-STAT3 signaling axis as a therapeutic target in breast cancer metastasis.

**One sentence summary:** LMO2 modulates STAT3 signaling in breast cancer metastasis.

## INTRODUCTION

While significant progress has been made to treat early-stage breast cancer, treatment options and outcomes for metastatic breast cancer have been largely unchanged in a decade (Esposito et al., 2021; Siegel et al., 2011; Siegel et al., 2021). In order to improve outcomes for breast cancer patients, it is critical to identify and elucidate signaling pathways active in metastatic cells. However, it has been difficult to pinpoint cancer cell populations involved in metastasis as they represent a transient state (Lu and Kang, 2019). Previous studies employing lineage tracing and cell surface marker profiling have implicated distinct subsets of tumor epithelial cells in breast cancer metastasis, primarily using lineage markers such as E-cadherin (Beerling et al., 2016, Padmanaban et al., 2019), N-cadherin (Li et al., 2020) and S100a4 (Fischer et al., 2015). Recent studies have also suggested that metastatic cells display hybrid features of both epithelial and mesenchymal lineages (Kröger et al., 2019; Pastushenko et al., 2021). This has led to a debate in the field about the precise molecular identity of metastasis-initiating cells (Lu and Kang, 2019; Shen and Kang, 2019; Ye et al., 2017).

Our previous work has demonstrated that in breast cancer, minority populations of phenotypically immature cells in the tumor are enriched in tumor-initiating potential and metastasis (Al-Hajj et al., 2003; Liu et al., 2010; Sikandar et al., 2017). Recent advances in single-cell technologies have revealed complex transcriptional landscapes in human tumors and enabled precise molecular characterization of these minority cell populations (Lawson et al., 2018). However, the functional and clinical significance of these populations remains to be elucidated (Lawson et al., 2018; Tanay and Regev, 2017). To understand the transcriptional heterogeneity in breast cancer, we performed single-cell RNA sequencing (scRNA-seq) in primary patient samples and developed a novel computational method that can predict immature cell populations *in silico* (Gulati et al., 2020). Using our scRNA-seq data, bulk tumor expression deconvolution, lineage tracing, and functional assays, we have now identified a clinically relevant population of metastasis-initiating cells that express the hematopoietic transcription factor and T-cell oncogene, LMO2. Here, we mechanistically define the role of LMO2 in breast cancer metastasis by its association with tumor vasculature and identify LMO2 as a previously unknown regulator of STAT3 signaling in breast cancer.

## RESULTS

### LMO2 is expressed in a minority population of immature THY1^+^/VEGFA^+^ human breast cancer cells

To understand the substructure of the epithelial populations in breast cancer, we started by analyzing scRNA-seq profiles (Gulati et al., 2020) of human breast tumor epithelial cells from patients with triple-negative (*n* = 5) or estrogen receptor positive (ER^+^) breast cancer (*n* = 13). We identified a minority population of *THY1*^+^ cells that were largely restricted to the basal compartment, comprising 11% of all basal cells (**Fig. S1A, Table S1**). Moreover, within this subset, 33% of cells expressed *VEGFA* (**Fig. S1A**). We were struck by this combination since THY1^+^ cells are enriched in reconstitution potential in the normal mammary gland (Lobo et al., 2018) and tumorigenic potential in mouse tumors (Cho et al., 2008) and VEGFA is a pro-angiogenic factor linked to tumor growth and distant metastasis (Mercurio et al., 2005; Zhao et al., 2015). To determine whether *THY1*^+^/*VEGFA*^+^ cells represent a potential immature cell population, we applied CytoTRACE, a computational framework for predicting cellular differentiation status on the basis of single-cell transcriptional diversity (Gulati et al., 2020). We found that relative to other basal cells, *THY1*^+^/*VEGFA*^+^ cells are predicted to be significantly less differentiated, suggesting a role for this population in tumor growth or metastasis (**Fig. 1A**).

**Figure 1:**
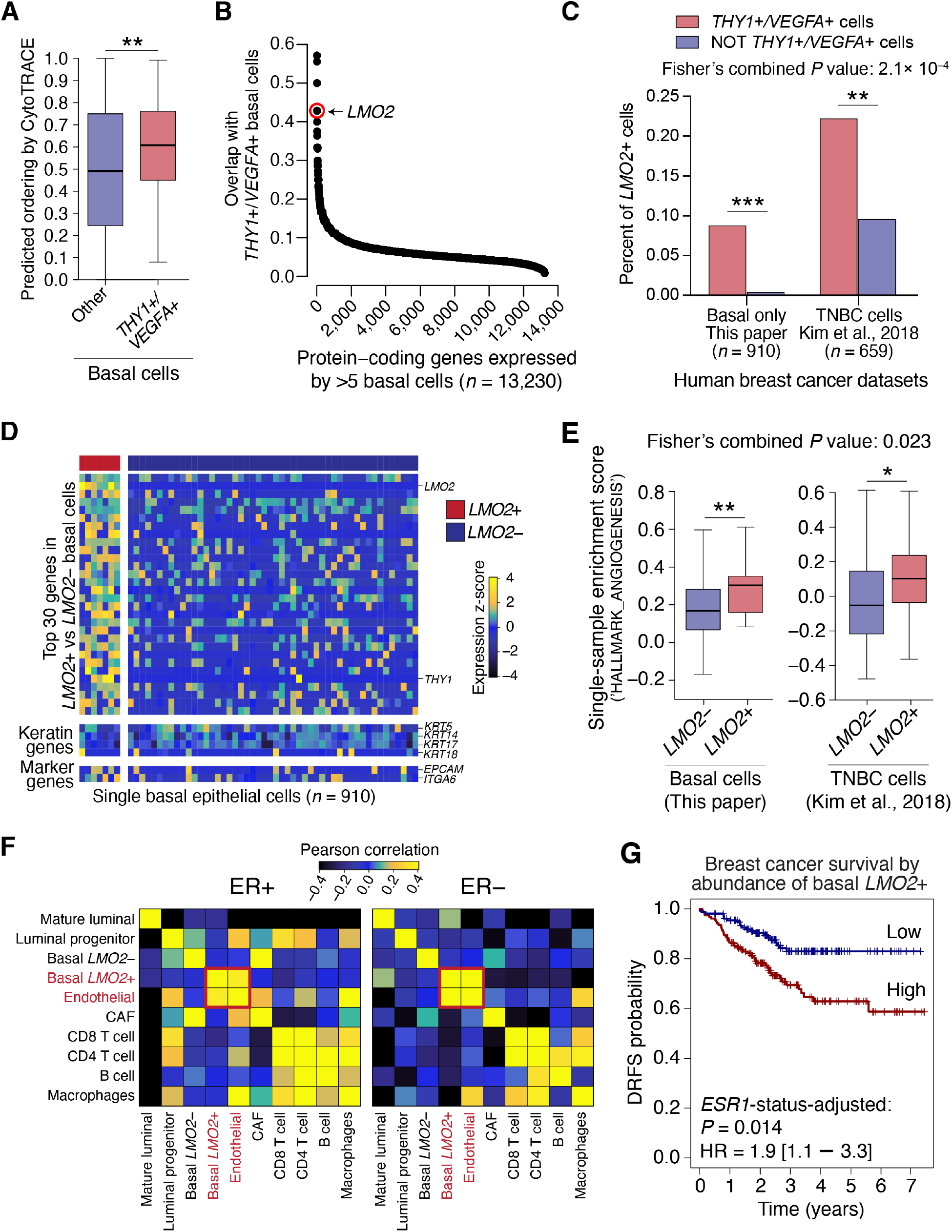
Identification of an immature basal epithelial population associated with pro-angiogenic signaling and poor survival in human breast cancer. (**A**) Differentiation scores of basal epithelial cells from 17 human breast tumors profiled by scRNA-seq (all but ‘SU196’ contained basal cells). Differentiation scores were determined by CytoTRACE (Gulati et al., 2020). Statistical significance between *THY1*^+^/ *VEGFA*^+^ basal cells and other basal cells was calculated using an unpaired two-tailed *t*-test. \**P*<0.1; \*\**P*<0.05; \*\*\**P*<0.01. (**B**) Plot showing protein-coding genes ordered by their enrichment in *THY1*^+^/ *VEGFA*^+^ basal cells from human breast tumors profiled by scRNA-seq. Enrichment was defined as the number of *THY1*^+^/*VEGFA*^+^ basal cells expressing a given gene (TPM > 0) divided by the total number of cells expressing that gene. Only genes expressed by at least 5 basal cells were considered. *LMO2* is highlighted in red. (**C**) Paired bar plots showing percent of *LMO2*^+^ cells in *THY1*^+^/*VEGFA*^+^ cells (red) and all other cells (blue) in two human breast cancer datasets, including Kim et al., 2018 (4 primary triple-negative breast cancers, single nucleus RNA-sequencing, tumor only, *n* = 659) (Kim et al., 2018), and the basal cells (see methods for details; *n* = 910) from this study. Statistical analysis was performed by Fisher’s Exact Test for association of *LMO2*^+^ cells with *THY1*^+^/*VEGFA*^+^ cells. Individual and combined *P* values by Fisher’s method are shown in the graph. \**P*<0.1; \*\**P*<0.05; \*\*\**P*<0.01. (**D**) Heatmap depicting the top 30 differentially expressed genes, along with selected keratin and lineage markers, in *LMO2*^+^ (*n* = 7 cells) vs. *LMO2*^−^ (*n* = 903 cells) basal epithelial cells from primary breast tumors. A random subsample of 50 *LMO2*^−^ basal cell transcriptomes is shown for clarity. Color scale (above) represents *z*-score-normalized expression per gene. (**E**) Differential enrichment of the ‘HALLMARK_ANGIOGENESIS’ pathway in *LMO2*^+^ vs. ^−^ in two independent human breast cancer datasets described in **C**. To ensure a fair comparison between *LMO2* positive and negative populations, an empirical *P* value was calculated by comparing the mean enrichment in *LMO2*^+^ basal cells versus a size-matched collection of *LMO2*^−^ basal cells randomly sampled 10,000 times. A combined *P* value by Fisher’s method is also shown. \**P*<0.1; \*\**P*<0.05; \*\*\**P*<0.01. **(F and G**) Cell-type and survival association of *LMO2*^+^ basal cells across 508 bulk human breast tumor transcriptomes (Esserman et al., 2012) deconvolved using CIBERSORTx. (**F**) Co-association patterns among cell type abundance profiles in bulk breast tumors, as quantified by Pearson correlation. Basal *LMO2*^+^ cells and endothelial cells are highlighted. (**G**) Kaplan Meier curves showing differences in distant recurrence-free survival (DRFS) in 508 breast cancer patients stratified by the median abundance of *LMO2*^+^ basal epithelial cells. DRFS was modeled as a function of *LMO2*^+^ basal cell status and *ESR1* status (**Methods**). The adjusted log-rank *P* value and hazard ratio with 95% confidence interval for *LMO2*^+^ basal cell status is shown.

To identify potential molecular regulators within this population, we next searched for genes with expression patterns that overlap *THY1* and *VEGFA* expression in our dataset. Intriguingly, we found that *LMO2*, a hematopoietic stem cell regulator (Yamada et al., 1998) and T-cell oncogene (Larson, 1995), was among the top five hits (**Fig. 1B, Table S2**). *LMO2* also marked *THY1*^+^/*VEGFA*^+^ cells in an independent scRNA-seq atlas of triple-negative human breast tumors (Kim et al., 2018), corroborating this result (**Fig. 1C**). Analysis of the *LMO2*^*+*^ basal epithelial subset showed that these cells not only express *THY1* and epithelial cytokeratins (**Fig. 1D**), but also display a coherent gene expression program significantly enriched in angiogenesis genes, including *VEGFA* and *S100A4* (**Fig. 1E, Table S3**).

We next measured the relative abundance of distinct endothelial, immune, stromal, and epithelial populations in human breast tumors with respect to *LMO2*^+^ basal cells. As *LMO2* is expressed in myriad cell types, including immune, stromal, and endothelial cells, the expression of the gene is insufficient to distinguish cell types. Therefore, we defined unique transcriptional signatures for various niche and breast epithelial cells from our scRNA-seq data and utilized CIBERSORTx, a deconvolution approach, to calculate the cellular composition of bulk RNA admixtures from breast cancer clinical cohorts (Newman et al., 2019) (**Methods**). In line with our previous results, we observed a striking correlation between the abundance of *LMO2*^+^ basal cells and endothelial cell content imputed in 508 breast tumors (Esserman et al., 2012) (*r* = 0.45; *P* < 2 × 10^−16^; **Fig. 1F**).

### Human *LMO2*^+^ basal cells are associated with poor outcomes in breast cancer patients

Deconvolution of an additional 3,024 human breast tumors from three clinical cohorts (Curtis et al., 2012; TCGA, 2012) revealed that basal *LMO2*^+^ cells are more abundant in ‘Basal’ breast cancer subtypes which correlate with more aggressive breast cancers as compared to other PAM50 classes (Perou et al., 2000) (**Fig. S1B**). We also found a significant increase in basal *LMO2*^+^ cells with worsening clinical grade and stage of the tumor (**Fig. S1C, D**), suggesting that *LMO2*^*+*^ cells increase with tumor progression. Importantly, higher levels of *LMO2*^+^ basal cells were significantly associated with inferior distant recurrence-free survival (**Fig. 1G**), independent of estrogen receptor status. These data link the abundance of *LMO2*^+^ basal epithelial cells with more aggressive breast tumors and distant metastasis.

### Lmo2 lineage-traced cells have a higher propensity to metastasize

To experimentally verify our *in silico* findings, we began by employing the CreERT2 system (Rios et al., 2014; van Amerongen et al., 2012; Van Keymeulen et al., 2011) to delineate the fate of epithelial cells that have expressed *LMO2*^+^ in breast tumors. We obtained *Lmo2*^*CreERT2*^ mice (Forster, Drynan, Pannell, Rabbitts *in preparation*) and crossed them to *Rosa26*^*mTmG*^ reporter and *MMTV-PyMT* tumor mice to generate triple-transgenic *Lmo2*^*CreERT2*^*/Rosa26*^*mTmG*^*/MMTV-PyMT* mice, which we termed *Lmo2-PyMT* (**Fig. 2A**). *MMTV-PyMT* tumors are an aggressive luminal subtype of breast cancer (Herschkowitz et al., 2007) that metastasize to the lungs (Guy et al., 1992) and have been extensively used to explore the cellular underpinnings of breast cancer metastasis (Beerling et al., 2016; Fischer et al., 2015; Padmanaban et al., 2019; Pastushenko et al., 2018). As *Lmo2* is expressed in other cells such as stromal and endothelial cells (Gratzinger et al., 2009), we orthotopically transplanted lineage depleted (CD45^−^/CD31^−^ /Ter119^−^) tumor cells from TdTomato-fluorescent *Lmo2*-*PyMT* into non-fluorescent BL6 mice to clearly assess the contribution of *Lmo2* lineage-traced breast cancer cells from the tumor. After tumors were formed, we pulsed the mice with tamoxifen to induce expression of GFP in *Lmo2*-expressing cells (**Fig. 2B**). At 48h post-pulse, we verified that expression of *Lmo2* was enriched in the transplanted GFP^+^ cancer cells (**Figs. 2C, S2**, and **S3A**). FACS quantification demonstrated that GFP^+^ cells represented a minor fraction of all tumor cells and expressed the epithelial marker, EpCAM (**Fig. 2C**).

**Figure 2:**
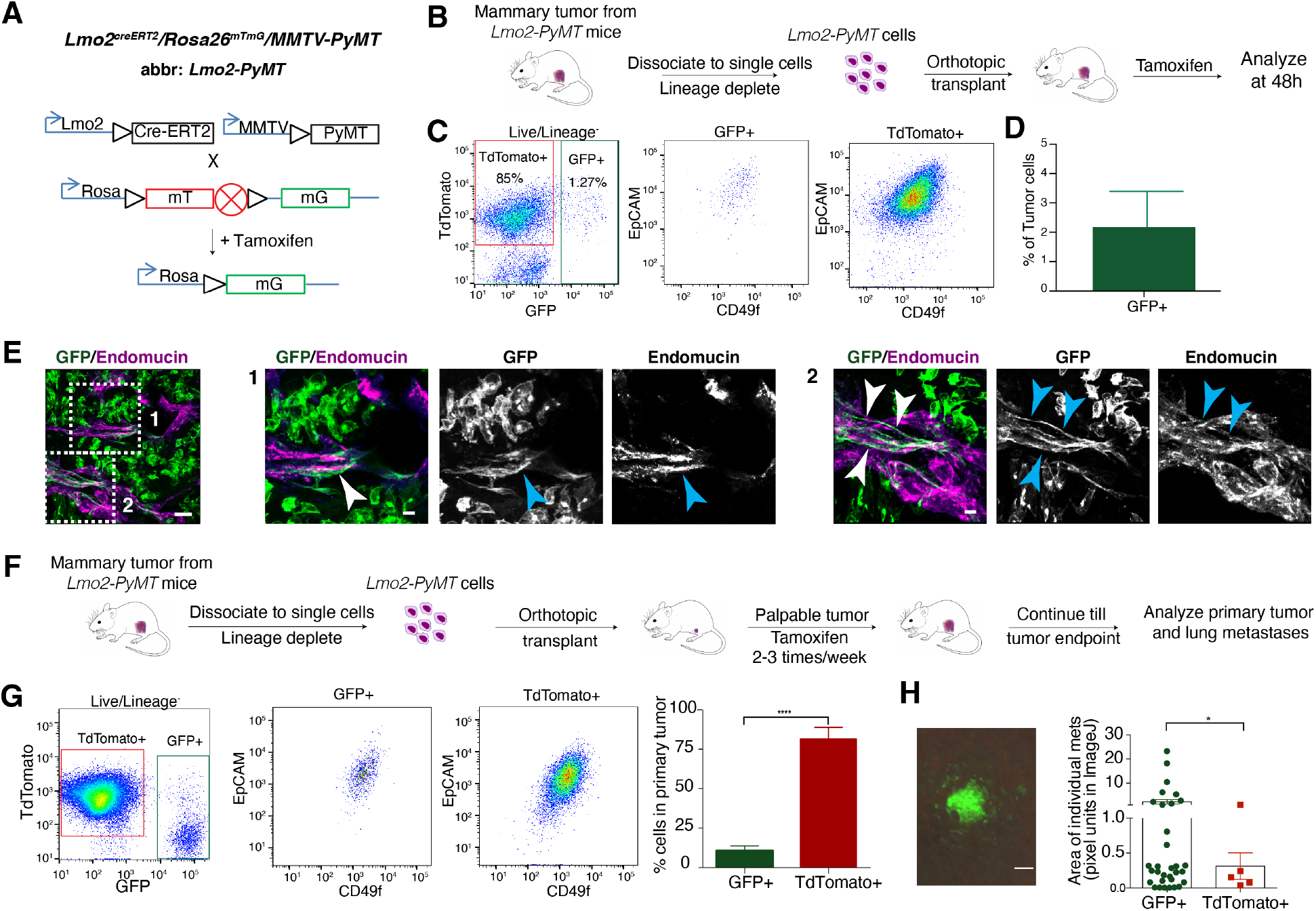
*Lmo2* lineage-traced tumor epithelial cells integrate into the vasculature and can form metastasis in *PyMT* tumors. (**A**) Schematic diagram showing generation of the triple transgenic *Rosa26*^*mTmG*^ reporter with *MMTV-PyMT* and *Lmo2-CreERT2* mice (referred to as *Lmo2-PyMT*). (**B**) Schematic diagram showing the experimental scheme for *Lmo2-PyMT* tumors treated with tamoxifen. (**C**) *Panel 1*: FACS analysis of *Lmo2-PyMT* tumors 48h after Tamoxifen pulse. Cells are gated on lineage^−^ (CD45^−^, CD31^−^, Ter119^−^), DAPI^−^ cells (See **Fig. S2**) and analyzed using TdTomato^+^ and GFP^+^. *Panels 2 and 3*: EpCAM and CD49f expression status in GFP^+^ and TdTomato^+^ cells. (**D**) Quantification of GFP^+^ cells from *Lmo2-PyMT* tumors (*n*=5 mice). (**E**) Representative immunofluorescence image of *Lmo2* lineage-traced cells (GFP^+^ green) co-localizing and integrating with endomucin (magenta) stained tumor vasculature. High resolution magnification of Inset 1 and 2 are presented, Scale bar = 50μm. (**F**) Schematic diagram showing the experimental scheme for *Lmo2-PyMT* tumors treated with tamoxifen to trace metastatic cells. (**G**) *Panel 1*: FACS analysis of *Lmo2-PyMT* tumors at tumor end point from (**F**). Cells are gated on lineage^−^ (CD45^−^, CD31^−^, Ter119^−^), DAPI^−^ cells (See **Fig. S2**) and analyzed using TdTomato^+^ and GFP^+^. *Panels 2 and 3*: EpCAM and CD49f expression status in GFP^+^ and TdTomato^+^ cells. *Panels 4:* Quantification of TdTomato^+^ and GFP^+^ cells from *Lmo2-PyMT* tumors (*n*=4 mice). (**H**) *Panel 1*: Representative image of metastasis shown, Scale bar = 100μm. *Panel 2*: Quantification of total number and area of GFP^+^ and TdTomato^+^ lung metastasis in *Lmo2-PyMT* tumors. (*n*=4 mice) Data are shown as mean ± SD, and statistical analysis was performed by unpaired, two-sided Wilcoxon rank sum test * *P*<0.05.

To assess the population dynamics of *Lmo2* lineage-traced cells, we plated TdTomato^+^ tumor cells from *Lmo2-PyMT* mice in 3D organoid assays and pulsed the organoids with 4-hydroxytamoxifen. Consistent with the *in vivo* model, lineage-traced GFP^+^ cells comprised a minority of tumor organoids (∼2%) 7 days post-pulse. This percentage was unchanged even after 4 weeks in culture, suggesting similar proliferative capacity between GFP^+^ and TdTomato^+^ cells (**Fig. S3B**). We confirmed this by plating sorted GFP^+^ and TdTomato^+^ cells in 3D organoid cultures and showing that both populations formed organoids at similar frequencies (**Fig. S3C**).

To determine whether *Lmo2*^*+*^ cells co-associate with endothelial cells, as predicted *in silico* (**Fig. 1F**), we stained vasculature with endomucin and visualized their co-localization with 3D imaging. We found that *Lmo2* lineage-traced cells not only resided near tumor blood vessels (**Fig. 2E**) but surprisingly ∼20% showed co-localization with tumor vasculature and appeared to be incorporated into the tumor vasculature (**Fig. 2E** and **S3D**).

Given that abundance of *LMO2*^+^ cells in patients predicts distant recurrence-free survival (**Fig. 1G**) and *Lmo2* lineage-traced cells reside closer to tumor vasculature, we next tested whether *Lmo2*^+^ cells have metastatic capabilities. As dissemination of metastatic cells occurs continuously during tumor growth, to lineage-trace tumor cells expressing *Lmo2*, we pulsed *Lmo2-PyMT* mice with tamoxifen 2-3 times per week once the tumors were palpable and continued until tumor endpoint **(see Methods**; **Fig. 3F**). At the end of the experiment, we found that in the primary tumor only 10-15% of tumor cells were GFP^+^ (**Fig. 2G**). Surprisingly, even though the tumor was majority TdTomato^+^, the lungs had a disproportionately higher number of GFP^+^ metastases, several of which were also larger than the TdTomato^+^ metastases (*P* = 0.034, Wilcoxon signed-rank unpaired test) (**Fig. 2H**). These data suggest that *Lmo2* lineage-traced cells have a higher propensity to form metastases in the *PyMT* mice and is consistent with our findings in human breast cancer patients (**Fig. 1G**). Furthermore, a subset of GFP tumor cells did not remain Lmo2 positive (**Fig. S3E**), suggesting that expression of Lmo2 in some cells represents a transient state, in agreement with previous studies linking transient cell states to metastases (Pastushenko et al., 2018).

**Figure 3.**
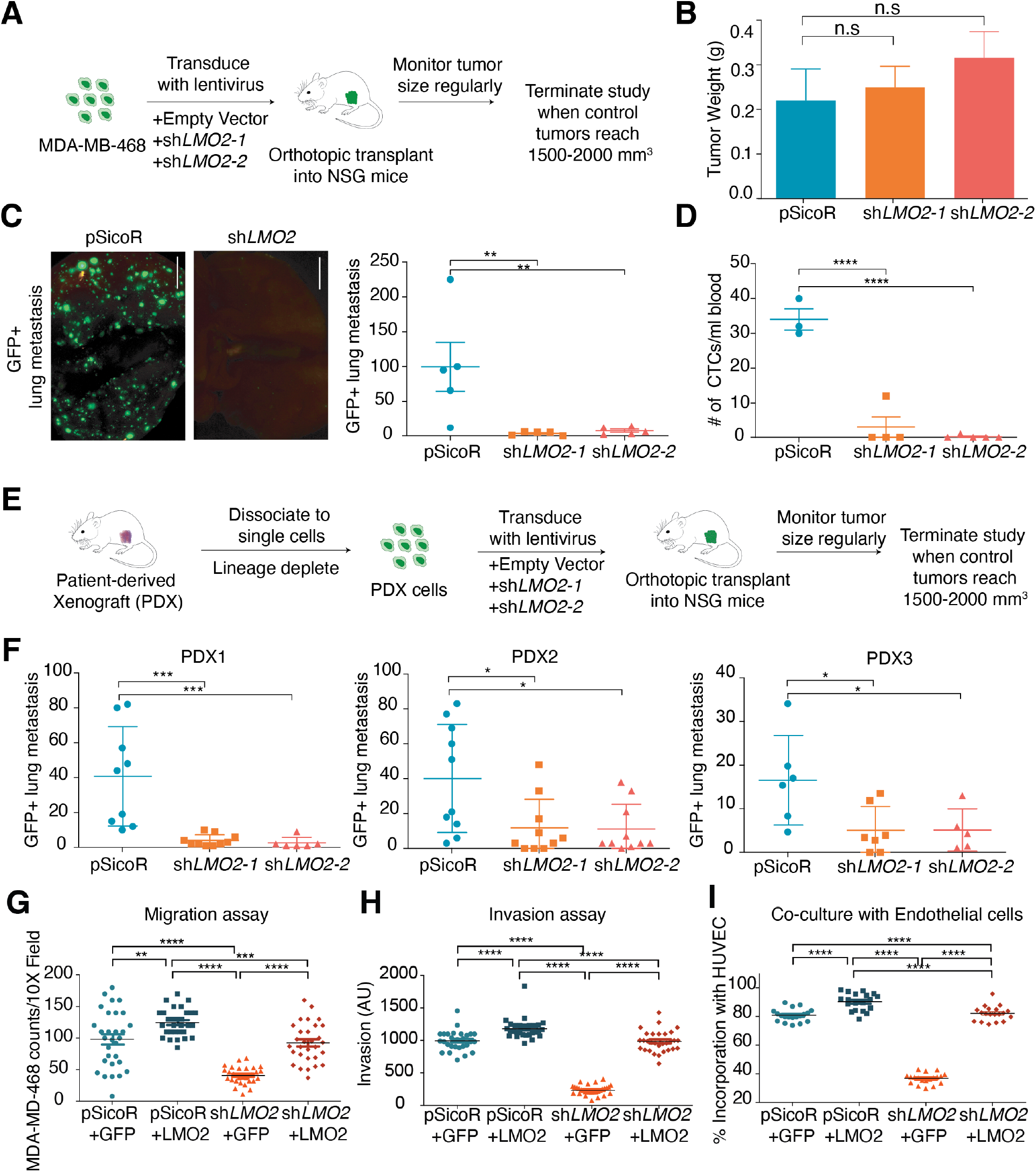
Knockdown of LMO2 reduces lung metastasis in human breast cancer. (**A**) Schematic of LMO2 knockdown in MDA-MB-468 cells followed by orthotopic transplant in NSG mice to evaluate tumor burden and metastases. (**B**) LMO2 knockdown in MDA-MB-468 cells. Tumor weight is shown with no significant difference between the control and LMO2 knockdown (*n*=5 mice/group). Data are shown as mean ± SD, and statistical analysis was performed by ANOVA with Dunnett’s adjustment, n.s *P*>0.05 (**C**) LMO2 knockdown decreases the number of spontaneous GFP^+^ lung metastasis in MDA-MB-468 cells (*n*=5mice/group). *Left panel*: representative immunofluorescence image with scale bar = 5mm, *right panel*: quantification. Data are shown as mean ± SD, and statistical analysis was performed by ANOVA with Dunnett’s adjustment, ** *P*<0.01. (**D**) LMO2 knockdown decreases the number of circulating tumor cells in MDA-MB-468 cells (*n*=3mice in pSicoR, 4 in sh*LMO2-1*, 5 in sh*LMO2-2*). Data are shown as mean ± SD, and statistical analysis was performed by ANOVA with Dunnett’s adjustment, **** *P*<0.0001. (**E**) Schematic of LMO2 knockdown in patient derived xenografts (PDXs) followed by orthotopic transplant in NSG mice to evaluate tumor burden and metastases. (**F**) LMO2 knockdown decreased number of spontaneous GFP^+^ lung metastasis in PDX samples. Data are combined from 3 independent experiments for PDX1, PDX3 and from 2 independent experiment for PDX2 (*n*=9 mice/ group for PDX1, *n*=6 mice/group for PDX2, *n*=10mice/group for PDX3). Data are shown as mean ± SD, and statistical analysis was performed by ANOVA with Dunnett’s adjustment, * *P*<0.05, ** *P*<0.01, *** *P*<0.001, **** *P*<0.0001. (**G**) MDA-MB-468 cells infected with shRNA targeting 3’ UTR of LMO2 or a control shRNA pSicoR were infected with either an empty vector control ‘GFP’ or an LMO2-overexpression vector ‘+LMO2’ to generate pSicoR +GFP, pSicoR +LMO2, shLMO2 +GFP, shLMO2 +LMO2. Transwell migration was quantification at 24 hours. (**H**) Spheroid invasion assay was performed and quantified at Day 5 using the breast cancer cells from (**G**). (**I**) The breast cancer cells from (**G**) were co-cultured with HUVEC cells and the percentage of breast cancer cells that are co-localizing with HUVEC tubes was quantified using ImageJ. For all experiments in (**G-I**), *n*=3 and 10 images were analyzed per condition per *n*. Statistical analysis was performed by ANOVA with Dunnett’s adjustment, and significance is indicated as ** *P*<0.01, *** *P*<0.001, **** *P*<0.0001

### LMO2 knockdown abrogates lung metastasis in human breast cancer models

To understand the functional role of LMO2 in human breast cancer, we knocked down *LMO2* expression in MDA-MB-468 cells using two independent shRNA vectors tagged with a GFP reporter (**Fig. S4A-C**). We then implanted the cells orthotopically in immunodeficient mice (**Fig. 3A** and **S4D**). In contrast to a previous report (Liu et al., 2016), knockdown of *LMO2* did not affect primary tumor growth (**Fig. 3B**) or proliferation *in vitro* (**Fig. S5A**). Nevertheless, *LMO2*-knockdown tumors had significantly fewer lung metastases relative to control (*P* = 0.003, ANOVA; **Fig. 2C**). Moreover, *LMO2*-knockdown in tumor-bearing mice led to a significantly reduced number of circulating tumor cells compared to control mice (*P* < 0.0001, ANOVA; **Fig. 2D**), implicating LMO2 in tumor cell shedding, a key step in metastasis initiation. To extend our findings to more clinically relevant models, we used patient-derived xenograft (PDX) models previously generated in our lab (Sikandar et al., 2017). Consistent with our MDA-MB-468 studies, knockdown of LMO2 dramatically decreased metastasis to the lung in three different PDX models of breast cancer (**Fig. 3E, F**), but did not significantly impact tumor growth (**Fig. S5B-D**).

To better understand how LMO2 affects metastasis, we rigorously studied the effects of LMO2 knockdown *in vitro* in MDA-MB-468 cells. Knockdown of LMO2 showed significant impairment in the ability of cancer cells to migrate across transwells and invade through a 3D hydrogel matrix (**Fig. S6A, B**). Importantly, since *LMO2*^+^ epithelial cells associated with endothelial cells in patient samples, we tested whether knockdown of *LMO2* decreased this association in co-culture assays. We found that in 3D co-culture assays with human vascular endothelial cells (HUVECs), LMO2 knockdown significantly impacted incorporation of cancer cells into HUVEC tubes (**Fig. S6C**). To confirm that the effects of knockdown were specific to LMO2, we overexpressed LMO2 in cells with shRNA targeting the 3’UTR. We found that all phenotypes of migration (**Fig. 3G**), invasion (**Fig. 3H**), and incorporation into the vasculature *in vitro* (**Fig. 3I**) could be rescued by overexpression of LMO2 in LMO2-deficient cells. Lastly, to test whether LMO2 is required after metastatic cells enter circulation, we injected control and LMO2 knockdown cells into the tail vein. We found that LMO2 knockdown did not significantly impact the formation of lung metastases when cells were directly injected in the tail vein, suggesting that LMO2 is critical for the initial dissemination of cancer cells from the tumor, but not extravasation and formation of metastatic foci (**Fig. S6D**).

### RNA sequencing identifies LMO2 as a regulator of IL6-JAK-STAT3 signaling

To elucidate the molecular function of LMO2 in breast cancer cells, we performed bulk RNA sequencing of MDA-MB-468 cells after transfection with control and LMO2 shRNA vectors (**Fig. 4A**). Among the top 50 genes downregulated after LMO2 knockdown were genes previously implicated in metastasis, such as *BMP2* (Bach et al., 2018; Huang et al., 2017; Wang et al., 2017), *LGR6* (Leushacke and Barker, 2012; Ruan et al., 2019), *EGR4* (Matsuo et al., 2014), *TDO2* (D’Amato et al., 2015) and *S100A4* (Boye and Maelandsmo, 2010; Garrett et al., 2006; Helfman et al., 2005) (**Fig. 4A, Table S4**). Using Gene Set Enrichment Analysis (GSEA) (Mootha et al., 2003; Subramanian et al., 2005), we found that inflammatory pathways, such as TNFα via NF-kB signaling, IL6-JAK-STAT3 signaling, and IFNγ response, were significantly downregulated in LMO2 knockdown as compared to control conditions (**Fig. 4B**). To confirm our findings in primary patient samples we performed single-sample GSEA in our scRNA-seq data set as well as a larger published dataset of primary human breast cancer cells (Kim et al., 2018). We found that IL6-JAK-STAT3 signaling was significantly enriched in *LMO2*^+^ versus *LMO2*^−^ single cells (**Fig. 4C**) compared to other pathways (**Fig. S7**). In the hematopoietic system, LMO2 is an adaptor protein that facilitates formation of functional protein complexes which then activate transcription of downstream targets (Chambers and Rabbitts, 2015). Hence, we asked whether LMO2 may similarly behave as a bridging molecule to drive downstream signaling in breast epithelial cells. Using proximity ligation assays, we found that LMO2 had a significantly high binding affinity to STAT3, but not to NF-kB, further confirming our pathway analysis (**Fig. 4D**).

**Figure 4:**
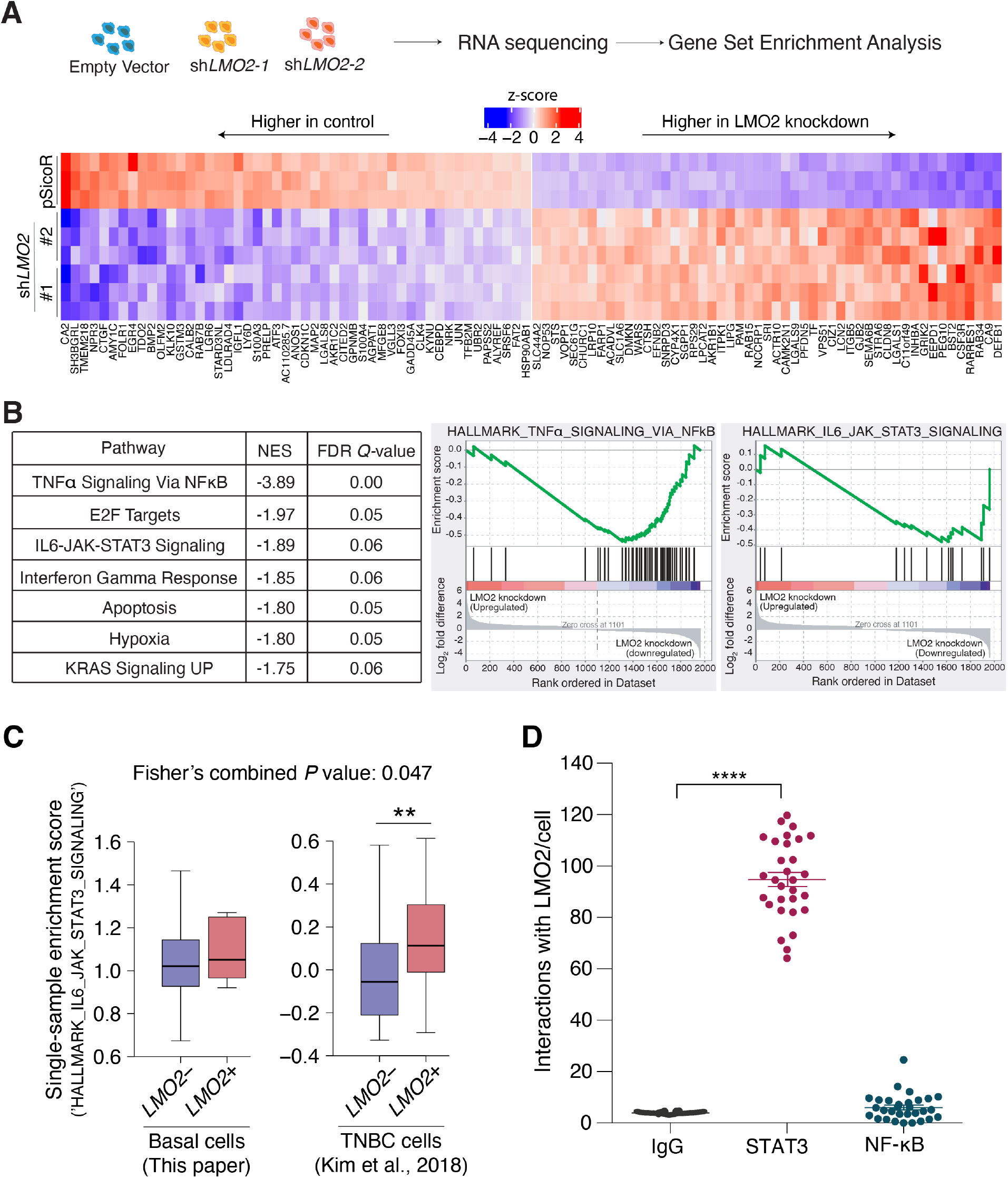
LMO2 regulates the IL6-JAK-STAT3 pathway and binds to STAT3. (**A**) *Top:* Schematic of bulk RNA-sequencing analysis in MDA-MB-468 cells infected with shRNAs targeting LMO2 or a control pSicoR. *Bottom*: Heatmap showing top and bottom 50 genes differentially expressed between control and LMO2 knockdown conditions, ordered by *P*-adjusted value. (**B**) *Left*: Hallmark gene sets found to be significantly enriched by GSEA analysis. Normalized enrichment scores (corresponding to control pSicoR vs LMO2 knockdown) and FDR *Q*-values are determined by the GSEA software. An FDR *Q*-value cutoff of <0.25 was used to select significant gene sets. *Right*: Enrichment plots for ‘HALLMARK_TNFα_SIGNALING_VIA _NFkB’ and ‘HALLMARK_IL6_JAK_STAT3_SIGNALING’ are depicted. (**C**) Differential enrichment of the ‘HALLMARK_IL6_JAK_STAT3_SIGNALING’ pathway in *LMO2*^+^ vs. ^−^ cells from two independent human breast cancer datasets as described in **Fig. 1C**. (**D**) Proximity mediated ligation assay showed that LMO2 had a stronger interaction with STAT3 compared to NF-kB *in vitro* (*n*=3, 10 images were analyzed per condition per *n*). Statistical analysis was performed by ANOVA with Dunnett’s adjustment. **** *P*<0.0001

### LMO2 is required for STAT3 activation by IL6 and TNFα

To demonstrate specificity and functional significance of the LMO2-STAT3 interaction, we first showed that LMO2 knockdown significantly reduced LMO2-STAT3 binding (*P* < 0.0001, ANOVA; **Fig. 5A**). We also confirmed the LMO2-STAT3 interaction using co-immunoprecipitation assays (Co-IP) of LMO2 with STAT3 (**Fig. 5B**) and, a reverse Co-IP of STAT3 with LMO2 (**Fig. 5C**). In breast cancer, STAT3 is activated by cytokines, such as IL6 (Zhong et al., 1994), TNFα (De Simone et al., 2015), IFNα (Beadling et al., 1994; Cho et al., 1996; Darnell et al., 1994) and IFNγ (Darnell et al., 1994; Will et al., 1996), as well as receptor tyrosine kinases such as EGFR (Kim et al., 2012; Zhao et al., 2020), leading to phosphorylation of STAT3. Dimerization of pSTAT3 and translocation to the nucleus activates transcription of downstream target genes involved in several processes, including metastasis. To understand whether the STAT3-LMO2 interaction has an effect on downstream STAT3 signaling, we used a STAT3-luciferase reporter assay. We stimulated control or LMO2 knockdown cells with IL6, TNFα, IFNγ, IFNα, and EGF. We found that cells with knockdown of LMO2 were unable to induce transcription of the STAT3-luciferase reporter when treated with IL6 and TNFα as compared to control (**Fig. 5D**), but STAT3-luciferase was activated by IFNγ, IFNα, and EGFR treatment. This suggests that LMO2 function in breast cancer cells is specific to activation of STAT3 signaling through IL6 and TNFα. On a molecular level, we found that knockdown of LMO2 significantly reduced STAT3 phosphorylation at Tyr705, which is required for its dimerization and transcriptional activity (**Fig. 5E and Fig. S8**). To understand how LMO2 regulates phosphorylation of STAT3, we examined the interaction of STAT3 with its upstream activator JAK2 and its cytoplasmic inhibitor PIAS3. Knockdown of LMO2 decreased the interaction of STAT3 with JAK2 (**Fig. 5F**) and allowed for increased interaction with its inhibitor, PIAS3 (**Fig. 5G**). This suggests that LMO2 works as an adaptor protein in the cytoplasm to stabilize the STAT3-JAK2 interaction, thereby allowing efficient phosphorylation and activation of STAT3 while simultaneously preventing its negative regulation by PIAS3 (**Fig. 5H**). This LMO2-mediated control of a core inflammatory response pathway could enable cancer cells to rapidly transition between cellular phenotypes required for metastasis and represents a therapeutic vulnerability that could be targeted.

**Figure 5:**
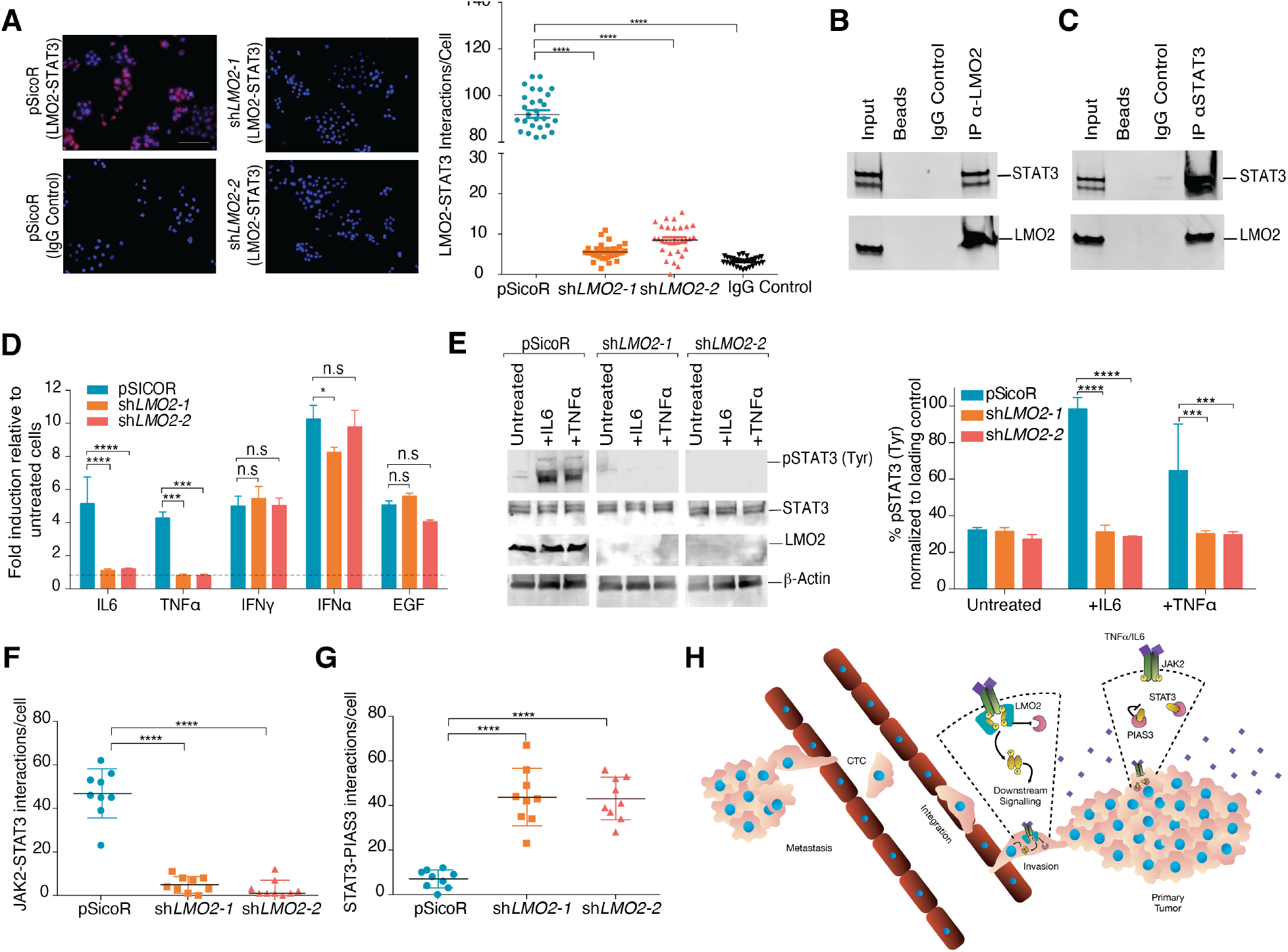
LMO2 stabilizes STAT3 signaling in breast cancer cells. (**A**) *Left panel*: Proximity mediated ligation assay shows that LMO2 binds to STAT3 *in vitro* and this interaction is significantly reduced with LMO2 knockdown indicating specificity of the assay. *Right panel*: Quantification of *n*=3 experiments and 10 images were analyzed per condition per *n*. Scale bar = 60μm. Statistical analysis was performed by ANOVA with Dunnett’s adjustment, **** *P*<0.0001. (**B**) Western blot of the input, immunoprecipitated beads (control), IgG (control) and LMO2 shows that LMO2 is able to pull-down STAT3. One representative blot of three independent experiments in shown. (**C**) Western blot of the input, immunoprecipitated beads (control), IgG (control) and STAT3 shows that STAT3 is able to pull-down LMO2. One representative blot of three independent experiments in shown. (**D**) STAT3-luciferase reporter activity shows robust stimulation of luciferase in control but not in cells with LMO2 knockdown when treated with IL6 and TNFα. IFNα, IFNγ, EGF treatment of cells results in robust stimulation in control and knockdown cells suggesting that LMO2 function is specific to IL6 and TNFα. Quantification of *n*=3 experiments. Statistical analysis was performed by 2-way ANOVA with Sidak’s correction, and significance is indicated as ** *P*<0.01, *** *P*<0.001, n.s. *P*>0.05. (**E**) Immunoblotting (*left panel*) and quantification (*right panel*) showed decreased phosphorylation of STAT3 at Tyr705 in LMO2 knockdown cells when treated with IL6 and TNFα indicating that LMO2 knockdown disrupts phosphorylation of STAT3. Quantification of *n*=3 experiments. Statistical analysis was performed by 2-way ANOVA with Sidak’s correction, and significance is indicated as \*\*\**P*<0.001, **** *P*<0.0001. (**F**) Interactions between STAT3 and JAK2 detected by proximity mediated ligation assay is reduced upon LMO2 knockdown indicating that LMO2 facilitates binding of STAT3 and JAK2. Quantification of n=3 experiments. Statistical analysis was performed by ANOVA with Dunnett’s adjustment, and significance is indicated as **** *P*<0.0001. (**G**) Interactions between STAT3 and PIAS3 detected by Proximity mediated ligation assay is increased upon LMO2 knockdown indicating that LMO2 prevents binding of STAT3 and PIAS3. Quantification of *n*=3 experiments. Statistical analysis was performed by ANOVA with Dunnett’s adjustment, and significance is indicated as **** *P*<0.0001. (**H**) Schematic of proposed mechanism of LMO2 in breast cancer metastasis. Tumor cells that express LMO2 have stabilized STAT3 signaling in response to IL6 and TNFα from the microenvironment, allowing these cells to intravasate into the circulation by incorporating into the vasculature.

## DISCUSSION

Efficient metastasis of tumor cells requires transition from a proliferative to an invasive state and back to a proliferative state at a distant site (Beerling et al., 2016). Previous studies using mouse tumor models have demonstrated the requirement of a basal epithelial program in metastasis (Cheung et al., 2013; Padmanaban et al., 2019) and showed that hybrid epithelial-mesenchymal states (Beerling et al., 2016; Kröger et al., 2019; Nieto et al., 2016) in metastasis express angiogenic factors (Pastushenko et al., 2018). Here, we have identified a population of *THY1*^+^/*VEGFA*^+^ human basal epithelial cells with higher transcriptional diversity that is marked by transient expression of *LMO2*. Moreover, we demonstrate that *Lmo2* lineage-traced epithelial cells have a higher propensity to form lung metastases. Moreover, knockdown of LMO2 decreases lung metastasis in multiple tumor models of human breast cancer by affecting multiple steps during intravasation. It is important to note that only a subset of *Lmo2* lineage-traced cells show vascular phenotypes, suggesting specific epigenetic regulation that is activated in the presence of TNFα and IL6 from the microenvironment. Our observations highlight a heterogenous, cancer-cell-intrinsic response to the microenvironment while previous studies have demonstrated that there is a reciprocal effect of cancer cells on the tumor microenvironment with recruitment of macrophages and cross-talk with tumor endothelial cells during metastasis (Borriello et al., 2020).

LMO2 has been extensively studied in hematological malignancies and is well-established as a transcriptional adaptor protein (Chambers and Rabbitts, 2015). Recent studies have attempted to understand the role of LMO2 in breast cancer (Hu et al., 2021; Liu et al., 2016; Liu et al., 2017) but have suffered from contradictory results, were limited to cell lines, and did not attribute LMO2 to any particular tumor cell population. We demonstrate that LMO2 is a previously unidentified binding partner of STAT3 in breast cancer cells and modulates STAT3 signaling in response to IL6 and TNFα. We speculate that the expression of LMO2 provides the necessary threshold to stabilize STAT3 signaling, which in turn enables the tumor cells to enter a transient metastatic state (Wendt et al., 2014) and escape the primary tumor. STAT3 signaling is involved in a number of processes and its targets may be defined in unison with other contextual signals such as inflammation. Several studies have linked low chronic inflammation in cancer to metastasis (Joyce and Pollard, 2009; Liu et al., 2015). We speculate that LMO2 is a critical molecular link between these processes and define a novel function for LMO2 in breast cancer metastasis. The development of new methods targeting adaptor proteins (Wang et al., 2020) and small molecules that disrupt the LMO2-STAT3 axis (Milton-Harris et al., 2020) could provide novel therapeutic strategies to modulate STAT3 signaling and inhibit metastatic colonization in breast cancer.

## ACKNOWLEDGMENTS

We thank Patricia Lovelace, Catherine Carswell Crumpton, and other flow cytometry staff for their help and animal facility core members. The Wolverine Aria instrument was funded by NHI grant S10-1S10RR02933801. We thank Diane Heiser for technical assistance with the tail vein injections. We thank Margaret Cuadro for administrative assistance. We thank the Stanford Neuroscience Microscopy Service, supported by NIH grant NS069375.

## Funding

This work was supported by NIH/NCI (U01CA154209-01 and P01 CA139490-05), the Breast Cancer Research Foundation (to M.F.C.), the U.S. Department of Defense (W81XWH-11-1-0287 and W81XWH-13-1-0281 to M.F.C.; S.S.S., W81XWH-12-1-0020), the National Cancer Institute (A.M.N., R00CA187192-03; M.F.C., 5R01CA100225-09; G.S.G., PHS grant no. CA09302), the Stanford Bio-X Interdisciplinary Initiatives Seed Grants Program (IIP) (A.M.N., M.F.C.), the Virginia and D.K. Ludwig Fund for Cancer Research (A.M.N., M.F.C), the Stinehart-Reed Foundation (A.M.N.), the Stanford School of Medicine Dean’s Fellowship (J.A.), Stanford Bio-X Bowes Graduate Student Fellowship (G.S.G.), and the Stanford Medical Science Training Program (G.S.G.). K.R-H. is a New York Stem Cell Foundation – Robertson Investigator. K.R-H. is also supported by the NIH (R01-HL128503)

## AUTHOR CONTRIBUTIONS

S.S.S. and M.F.C. conceived and designed the study. S.S.S. and J.A. performed experiments and analyzed data with supervision from M.F.C. G.S.G. analyzed single-cell and bulk RNA sequencing data with assistance from C.B.S. and supervision from A.M.N. A.H.K. assisted with the PDX studies. W.H.D.H. assisted with the metastasis experiments. S.D. performed staining for visualization of tumor vasculature under the supervision of K.R-H. T.A.P. assisted with the circulating cells experiment under the supervision of P.B. D.Q. provided technical support. F.D. assisted with the collection of patient specimens. J.P.T. assessed the enrichment of genes in LMO2^+^ cells and provided guidance in the project. T.R provided the *Lmo2*^*CreERT2*^ mice. S.S.S., J.A., G.S.G., A.M.N. and M.F.C., wrote the manuscript. All authors commented on the manuscript.

## Supplementary Materials

Materials and Methods

Figures S1-S8

Tables S1-S5

## Notes

https://www.ncbi.nlm.nih.gov/geo/query/acc.cgi?acc=GSE159285

### Competing Interest Statement

A.M.N. has patent filings related to expression deconvolution and digital cytometry, and has served as a consultant for Roche, Merck and CiberMed. A.M.N., G.S.G., S.S.S., and M.F.C. have patent filings related to differentiation prediction and therapeutic targets for breast cancer.

