## Supplemental information for "LMO2 is critical for early metastatic events in breast cancer"

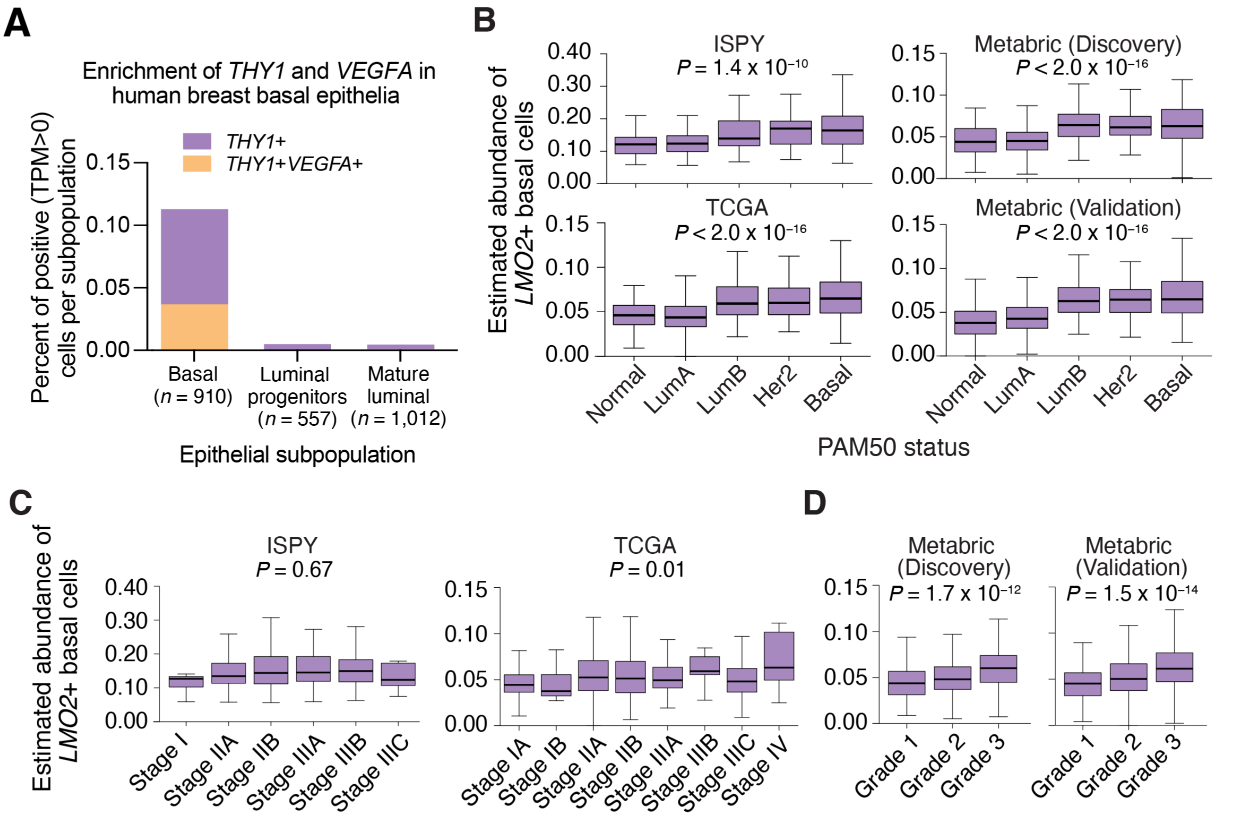


**Supplemental Figure 1: Expression of *THY1*, *VEGFA*, and *LMO2* in human breast epithelia and association of *LMO2*$^+$ basal cells with PAM50 subtype, clinical grade, and stage.**

(**A**) Stacked bar plots showing percent of cells expressing *THY1* (purple) and both *THY1* and *VEGFA* (orange) across three human breast epithelial subpopulations: basal (*n*=910), luminal progenitor (*n*=557), and mature luminal (*n*=1,012).

(**B to D**) Boxplots showing the estimated abundance of *LMO2*$^+$ basal cells in ISPY, TCGA, and Metabric (Discovery and Validation) by (**B**) PAM50 subtype, (**C**) stage, and (**D**) grade. *P* values were calculated by one-way ANOVA.

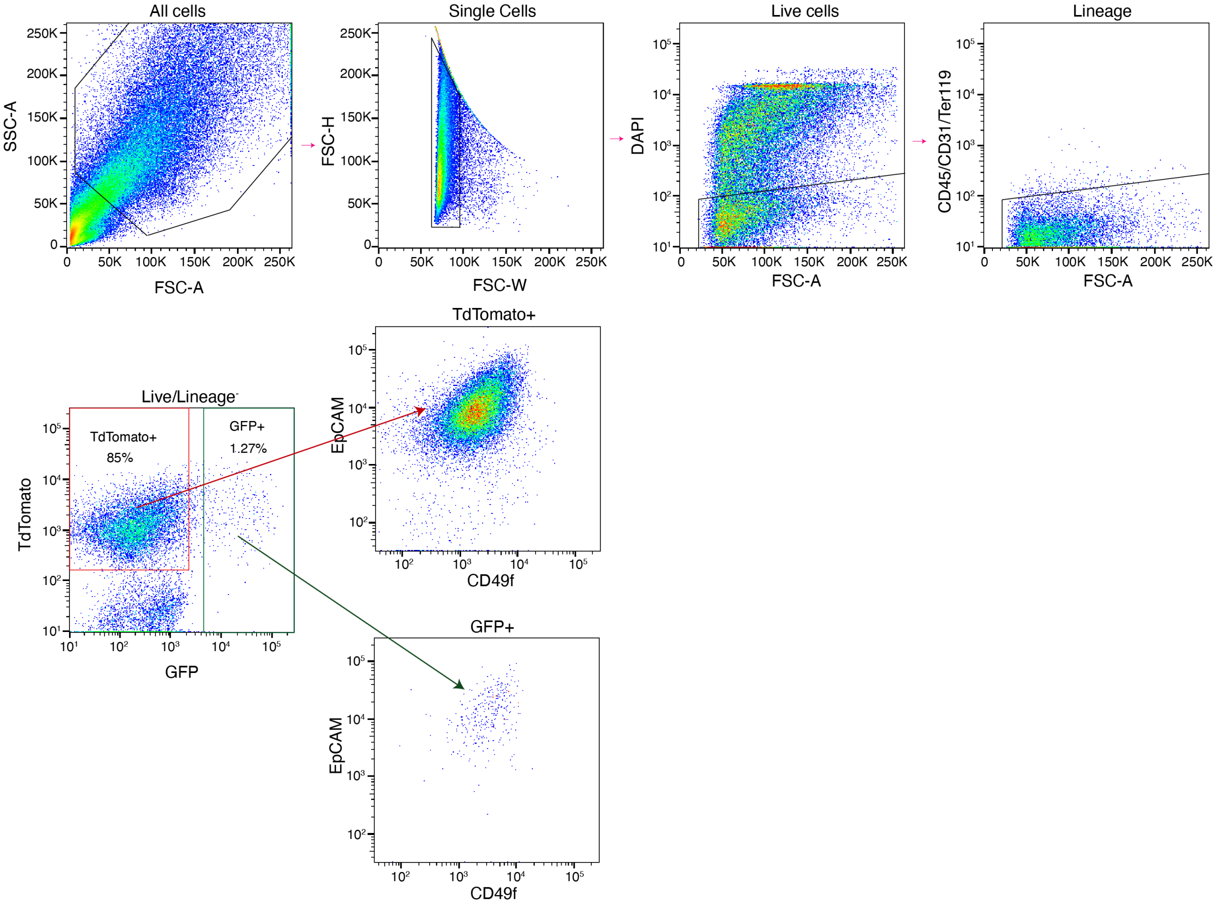


**Supplemental Figure 2: Gating scheme on *Lmo2-PyMT* mice.**
FACS analysis of *Lmo2-PyMT* tumors 48 hours after tamoxifen pulse in recipient non-fluorescent mice as shown in **Fig. 2B.** Cells are gated on lineage$^-$ (CD45$^-$, CD31$^-$, Ter119$^-$), DAPI$^-$. TdTomato$^+$ and GFP$^+$ are then analyzed using EpCAM and CD49f. *Lmo2* lineage-traced cells are EpCAM$^+$/CD49f$^+$.

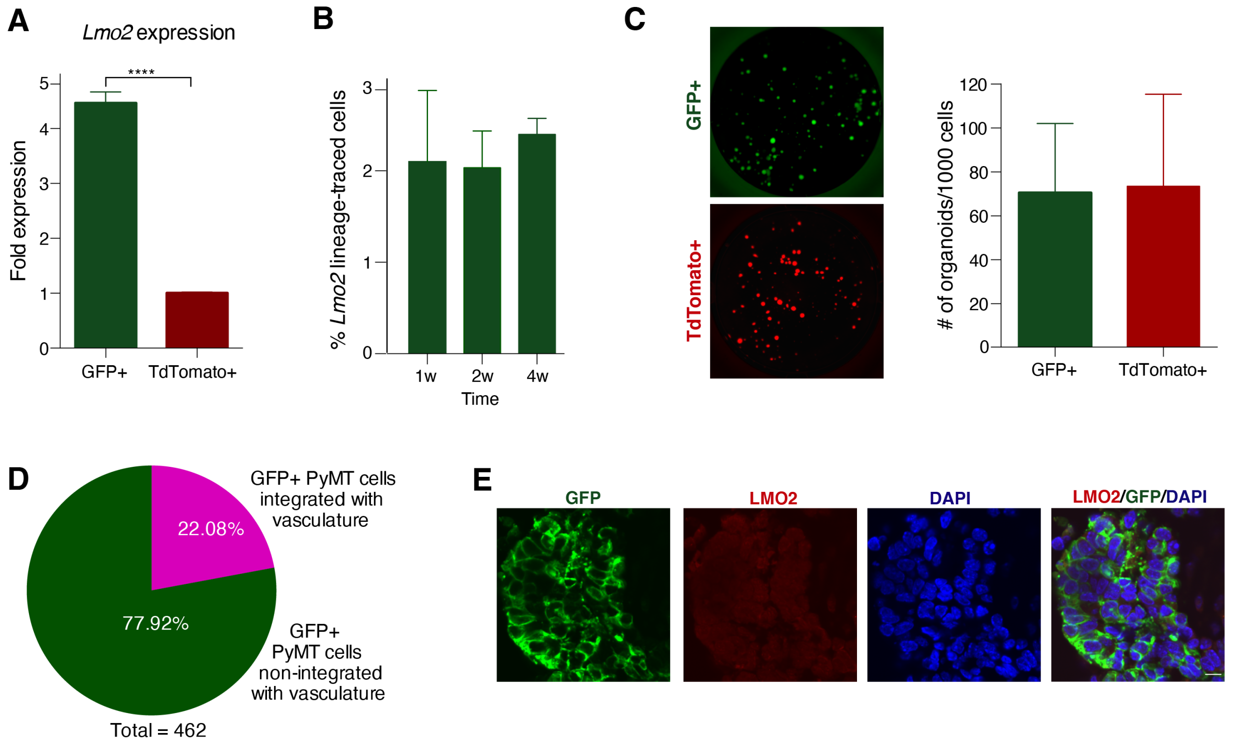


**Supplemental Figure 3: Population dynamics of *Lmo2* lineage-traced cells.**
(**A**) *LMO2* expression in TdTomato$^+$ and GFP$^+$ from *Lmo2-PyMT* tumors 48 hrs post tamoxifen pulse (*n*=4 mice). Student's *t*-test and *****P*<0.0001.
(**B**) Percentage of *Lmo2* lineage-traced cells in mammary organoids *in vitro* at 1-, 2- and 4-weeks after tamoxifen pulse (*n*=3 independent experiments performed in triplicates).
(**C**) Organoids formed by sorted TdTomato$^+$ or *Lmo2* lineage traced GFP$^+$ cells (Left panel: representative images, Right panel: quantification).
(**D**) Quantification of GFP$^+$ *PyMT* cells that integrate into the tumor vasculature from **Fig. 2E**.
(**E**) Representative immunofluorescence image of *Lmo2* lineage traced GFP$^+$ cells and LMO2 staining in *Lmo2-PyMT* tumors pulsed with tamoxifen indicating that *LMO2* expression is transient during metastasis, Scale bar=10μm.

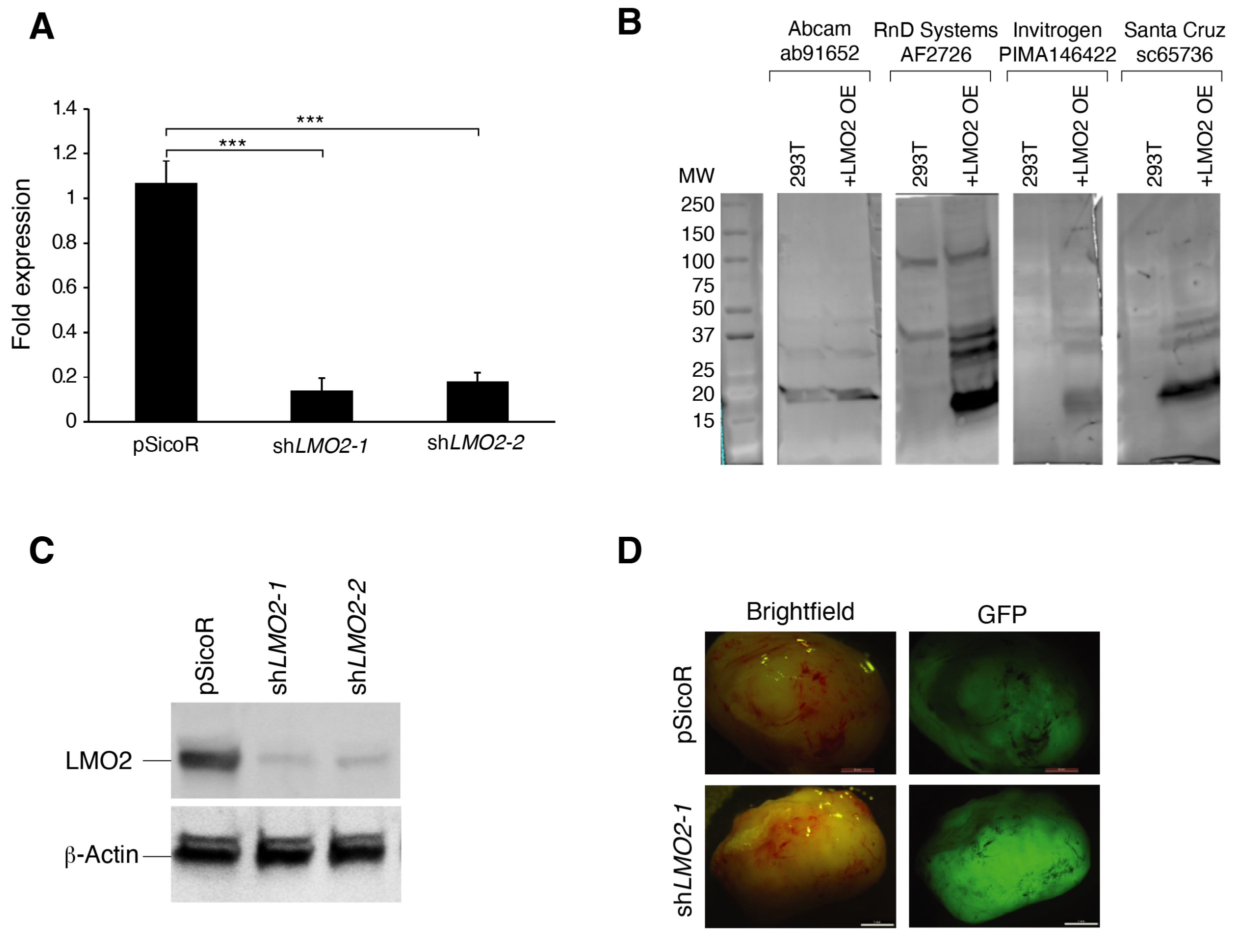


**Supplemental Figure 4: Knockdown of LMO2 in human breast cancer cells.**
(**A**) *LMO2* expression in MDA-MB-468 cells infected with pSicoR, sh*LMO2-1* and sh*LMO2-2* (*n*=4). Statistical analysis was performed by ANOVA with Dunnett's adjustment, and significance is indicated as *** *P*<0.001.
(**B**) Immunoblot for testing anti-LMO2 antibodies in 293T cells (null for LMO2 expression) and 293T that are infected with an LMO2 over-expression vector (293T+LMO2 OE). *Note: The abcam antibody (#91652) previously used to elucidate the binding partners and molecular role of LMO2 in breast cancer metastasis does not recognize LMO2 but instead detects another protein band at the same molecular weight.*
(**C**) Immunoblotting of LMO2 protein in MDA-MB-468 cells infected with shRNAs targeting LMO2 as compared to control pSicoR. β-Actin is used as a loading control.
(**D**) Representative immunofluorescence and brightfield images of MDA-MB-468 tumors formed in NSG mice by cells infected with a lentiviral vector with GFP as a reporter and pSicoR and sh*LMO2-1*.

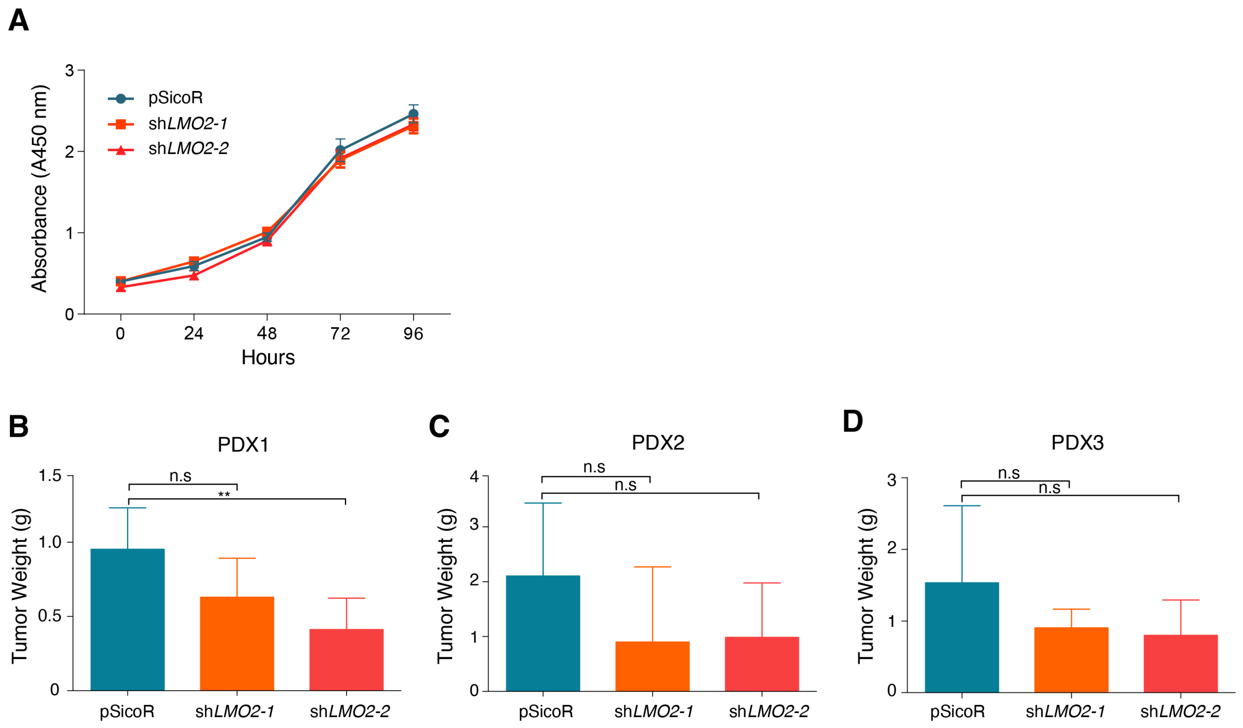


**Supplemental Figure 5: Effect of LMO2 knockdown in breast cancer cells proliferation.**
(**A**) Proliferation measured by WST1 absorbance between 0-96 hours in MDA-MB-468 cells infected with shRNAs targeting LMO2 as compared to control pSicoR (*n*=3).
(**B-D**) Tumor weights from 3 patient-derived xenografts (PDXs) infected with control pSicoR or shRNA targeting LMO2. Data are combined from 3 independent experiments for PDX1, PDX3, and from 2 independent experiment for PDX2 (*n*=9 mice/group for PDX1, *n*=6 mice/group for PDX2, *n*=10 mice/group for PDX3). Data are shown as mean ± SD. Statistical analysis was performed by ANOVA with Dunnett’s adjustment, and significance is indicated as $^{**}P < 0.01$; n.s $P > 0.05$.

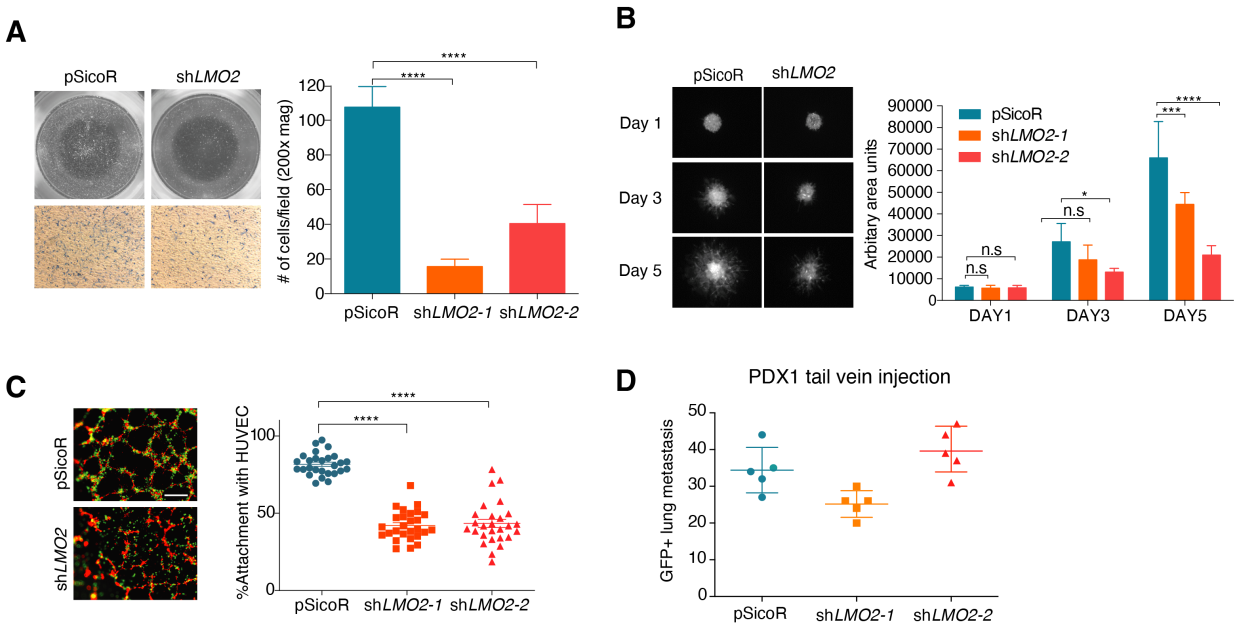


**Supplemental Figure 6: Effect of LMO2 knockdown in intravasation, migration, extravasation, and metastatic foci formation.**

*In vitro* properties of MDA-MB-468 cells upon LMO2 knockdown:

(**A**) Migration. *Left panel*: Representative image at *t*=36 hours. *Right panel*: Quantification of migration at 36 hours.

(**B**) Invasion. *Left panel*: Representative images at *t*=1, 3 and 5 days. *Right panel*: Quantification of invasion.

(**C**) Integration with HUVECs. *Left panel*: Representative images at 12 hours, Scale bar=100 mm; *Right panel*: Quantification of percentage of breast cancer cells that integrated with HUVEC tubes using ImageJ.

For all experiments, *n*=3 and 10 images were analyzed per condition per *n*. Statistical analysis was performed by ANOVA with Dunnett's adjustment, and significance is indicated as * *P*<0.05, ** *P*<0.01, *** *P*<0.001, **** *P*<0.0001.

(**D**) Lung metastasis after tail vein injection of PDX1 cells infected with control, sh*LMO2-1*, or sh*LMO2-2* vectors to assess the function of LMO2 in extravasation and metastatic foci formation.

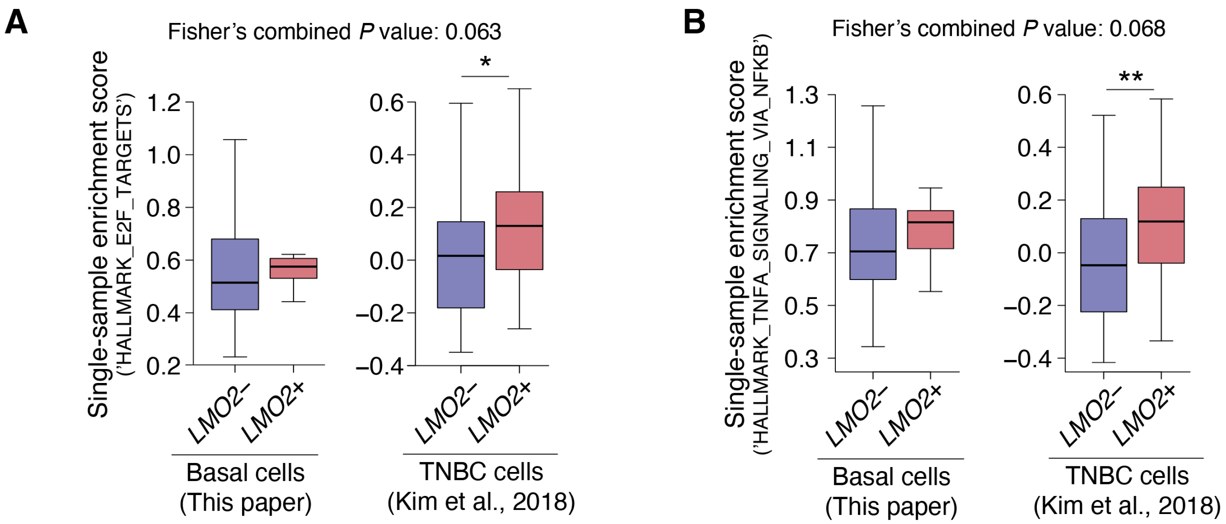


**Supplemental Figure 7: HALLMARK pathway enrichment in human breast cancer scRNA-seq datasets.**
(**A and B**) Differential enrichment of the (**A**) ‘HALLMARK_E2F_TARGETS’ and (**B**)‘HALLMARK_TNFA_SIGNALING_VIA_NFKB’ pathway in *LMO2*$^{+}$ vs. $^{-}$ cells from two independent human breast cancer datasets as described in **Fig. 1C.** *$P$<0.1; **$P$<0.05.

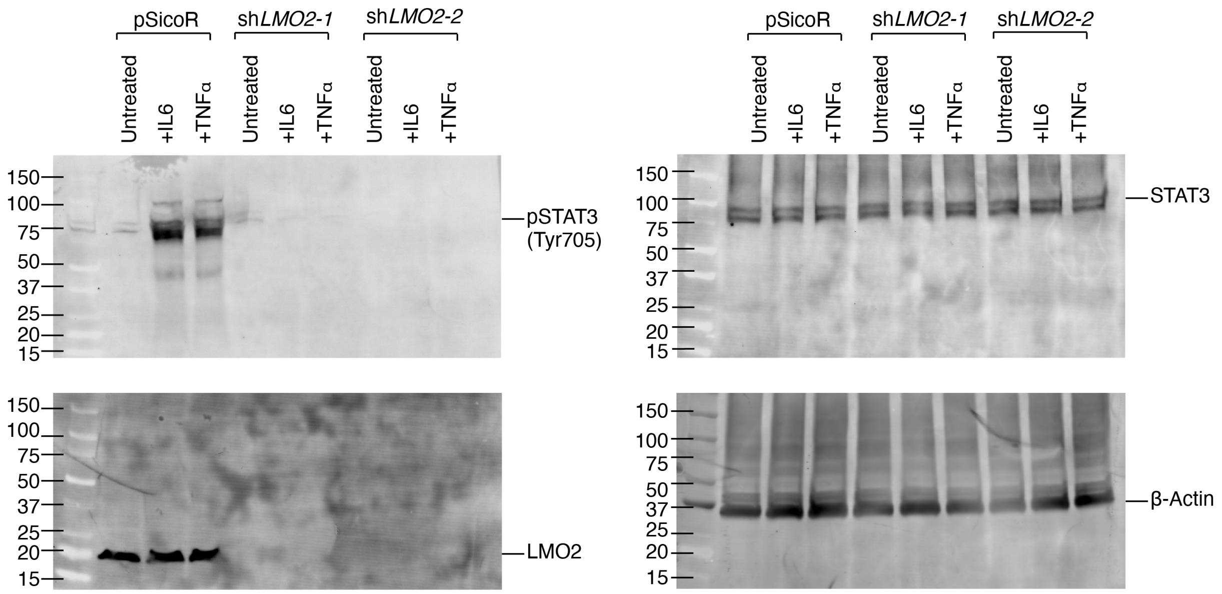


**Supplemental Figure 8: Full-length blots probed for pSTAT3, STAT3, LMO2 and β-Actin upon IL6 and TNF-α stimulation of MDA-MD-468 cells.**

### MATERIAL AND METHODS

**Preparation of single-cell suspensions for human and mouse tissues:**

For human samples, informed consent was obtained after the approval of protocols by the Stanford University and City of Hope Institutional Review Boards (IRB #4344). Tumor biopsies from human breast cancer patients (*n*=18) were obtained from the primary site (*n*=16), lymph nodes (*n*=1; paired primary), or brain metastasis (*n*=2) during surgical resection of breast tumors at Stanford Hospital and City of Hope National Medical Center (Table S1). Samples were mechanically dissociated into < 1-2 $mm^3$ pieces with a razor blade and then digested at $37^o$ C with 1,500 U collagenase and 500 U hyaluronidase in Advanced DMEM/F-12 (Thermo Fisher Scientific), 2 mM Glutamax (Invitrogen), and an antibiotic/antimycotic mix containing 120 µg/mL penicillin, 100 µg/mL streptomycin, and 0.25 µg/mL amphotericin-B (PSA) for 4-6 hrs with hourly pipetting for 5 min. After digestion, cells were treated with ACK lysis buffer to deplete red blood cells and then incubated with 10 U dispase to further dissociate the tissue into single cells and 1,000 U DNase I to prevent cell clumping. Cells were filtered through a 70 µm nylon mesh and washed with staining buffer containing 2% fetal bovine serum (FBS) and PSA in Hank's Balanced Salt Solution (HBBS). Single cell suspensions of fresh breast tissue were then stained with fluorescent antibodies to prepare for FACS.

**Flow cytometry:**

To reduce nonspecific antibody binding, single cells were blocked with 10 µg/mL rat IgG on ice for 10 mins. Cells were then stained, in the dark, on ice for 30 mins. FACS was performed with a 130 µm nozzle on a BD FACSAria II with BD FACSDiva software. Side scatter and forward scatter profiles (area and width) were used to eliminate debris and cell doublets. Dead cells were eliminated by excluding 4',6-diamidino-2-phenylindole (DAPI) positive cells. For single-cell RNA-sequencing (scRNA-seq), human breast epithelia and niche cells were isolated as described in the section below ('*Single cell RNA sequencing analysis*'). For mouse *Lmo2-PyMT* tumor studies, tumor epithelial cells were enriched by negative gating of lineage markers CD45, Ter119, and CD31. For xenograft studies, human tumor epithelial cells were enriched by negative gating of H-2kd, a surface protein expressed on all mouse cells. A complete list of antibodies is provided in Table S5.

**Single-cell RNA sequencing:**

1,902 scRNA-seq profiles of tumor and adjacent-normal human breast basal (*n*=660), luminal progenitors (*n*=532), and mature luminal (*n*=710) cells were acquired from a previous study (accession code: GSE138536; Gulati et al., 2020). Using the same strategies for cell sorting, library construction, and data processing as previously described (Gulati et al., 2020), 250 additional basal epithelia and 207 stromal cells were collected from human breast tumors. Briefly, by negative gating of lineage cells expressing CD45, CD31, CD3, CD16, and CD64, cancer-associated fibroblasts (CAFs; *n*=6) and endothelial cells (ECs; *n*=126) were sorted as Lineage$^{-}$CD49f$^{-}$EpCAM$^{-}$. Although ECs are generally CD31$^{+}$, the chemical digestion for tumor isolation cleaves surface CD31, so these cells were detected on flow cytometry as CD31$^{-}$. CAFs and ECs were then distinguished based on transcriptional expression of *FAP* and *PDGFRA* in CAFs and *CDH5, EMCN,* and *PECAM1* in ECs. Hematopoietic populations were sorted separately based on surface expression of CD45 followed by CD14$^{+}$ for macrophages (*n*=21), CD3$^{+}$CD4$^{+}$ for CD4 T cells (*n*=22), CD3$^{+}$CD8$^{+}$ for CD8 T cells (*n*=21), and CD19$^{+}$ for B cells (*n*=11). A matrix containing gene-level transcripts per million (TPM) for all single cells (*n*=2,359) has been deposited in Gene Expression Omnibus (GEO) under the accession GSE159285. Metadata for each single cell is also available in GSE159285 and in Table S1.

**Predicted ordering of single cells by differentiation status:**
Single-cell level prediction of differentiation states in human breast tumor basal epithelia (*n*=910) was performed in R using the CytoTRACE package publicly available at https://cytotrace.stanford.edu (Gulati et al., 2020).

**Deconvolution of bulk breast tumors:**
CIBERSORTx was used to deconvolve cell type abundances from (1) microarray gene expression data of 508 bulk breast tumors from the Investigation of Serial Studies to Predict Your Therapeutic Response with Imaging and Molecular Analysis (I-SPY1) clinical trial (Esserman et al., 2012); (2) microarray gene expression data of 1,981 bulk breast tumors from Metabric Discovery (*n*=995) and Validation (*n*=986) cohorts (Curtis et al., 2012); (3) RNA-seq data from 1,033 bulk breast tumors from The Cancer Genome Atlas (TCGA) (Koboldt et al., 2012). Default parameters as described in the 'Tutorial' page at http://cibersortx.stanford.edu/ were used to generate a signature matrix from scRNA-seq data. Quantile normalization was run on microarray (but not RNA-seq) data and bulk-mode batch correction (B-mode) was applied for cross-platform deconvolution.

**Single sample GSEA analysis:**
Single-sample enrichment of the 'HALLMARK_ANGIOGENESIS', 'HALLMARK_E2F_TARGETS', 'HALLMARK_TNFA_SIGNALING_VIA_NFKB', and 'HALLMARK_IL6_JAK_STAT3_SIGNALING' gene sets was calculated using single-sample gene set enrichment analysis (ssGSEA) as implemented in the R GSVA package (v1.30.0)(Hänzelmann et al., 2013).

**Survival analysis of *LMO2*$^{+}$ basal cells:**
The *survival* v3.1.12 R package was used to analyze the association of *LMO2*$^{+}$ basal cells with distant recurrence-free survival (DRFS) in the I-SPY1 cohort (*n*=508 human breast

cancer samples). Samples were evenly stratified into 'High' and 'Low' groups based on whether tumors had greater than or less than the median abundance of *LMO2*$^+$ basal cells, respectively. A Cox proportional hazard model was then used to calculate the effect of *LMO2*$^+$ basal cell abundance on DRFS, adjusting for *ESR1* status as a possible confounder (*DRFS ~ LMO2pos_basal_status* + *ESR1_status*).

**Mice:**

*Rosa26*$^{mTmG}$ (Stock #007576), C57BL6 (Stock #000664) and *NOD.Cg-Prkdc*$^{scid}$*Il2rg*$^{tm1Wjl}$*/SzJ* (NSG) (Stock #005557), *MMTV-PyMT* (Stock #022974), mice were purchased from the Jackson Laboratory. *Lmo2*$^{CreERT2}$ transgenic mice were generated by pro-nuclear injection in C57Bl/6/CBA (Forster, Drynan, Pannell, Rabbitts in preparation). All tumor xenotransplantation was done in NSG mice. All mice used for this study were maintained at Stanford Animal Facility in accordance with the guidelines of the animal care use committee (APLAC #10868).

**Cell lines:**

MDA-MB-231, MDA-MB-468, and 293T cells were obtained from ATCC. These cells were certified by the vendors to be mycoplasma free. None of the cell lines used are listed in the database of commonly misidentified cell lines maintained by ICLAC. Cell lines have not been authenticated but all cell lines used were passaged less than 10 times from when the original cells from the vendors were thawed. Cell lines are routinely tested for mycoplasma contamination and were mycoplasma free. MDA-MB-231, MDA-MB468, and 293T cells were grown in DMEM (Invitrogen) supplemented with PSA, 10% FBS (Hyclone), Glutamax (ThermoFisher Scientific), and sodium pyruvate (Life Technologies).

**Immunofluorescence staining in paraffin sections:**

Tumors were fixed in formalin and embedded in paraffin for immunostaining. Sections were de-paraffinized, dehydrated, and microwaved for 20 mins at 95 °C in Sodium Citrate Buffer (10mM Sodium Citrate, 0.05% Tween 20, pH 6.0) for antigen retrieval. Tissue sections were incubated overnight at 4 °C with primary antibodies diluted in phosphate buffered saline (PBS) + 5% Bovine Serum Albumin (BSA) (Antibodies are listed in Table S5). Samples were subsequently washed with PBS and were incubated with anti-GFP in Alexa Fluor 488 (1:500) and anti-mouse Alexa Fluor 594 conjugated secondary antibodies (Invitrogen) at 1:500 in PBS + 5% BSA for 1 hr at RT. All the immunofluorescence sections and cells were mounted in ProLong Gold with DAPI. Images were acquired by Carl Zeiss LSM 710 Meta confocal microscope. Images were processed using ImageJ.

**Real Time PCR:**

10,000 GFP$^+$ or TdTomato$^+$ *PyMT*-tumor cells were sorted into 1.5 microfuge tubes and spun down. RNA was extracted using the RNeasy Micro Kit (Qiagen #74034). RNA was reverse transcribed to cDNA using SuperScript III First Strand Synthesis kit (Life Technologies #11752-050) according to the manufacturer's instructions. cDNA was preamplified 15 cycles according to the cell number using TaqMan pre-amp mastermix (Applied Biosystems #4391128) and target gene Taqman primer pool. Preamplified cDNA was then subjected to the real time PCR for specific gene target according to manufacturer's instruction using 7900HT Real Time PCR system (Applied Biosystems). All expression data are normalized to *Actnb* and *Gapdh*. Data were analyzed by SDS2.4 software and Excel.

**Whole mount immunostaining in tumors:**

Procedure was performed as previously described (Das et al., 2019). Tumors were dissected in PBS, immediately fixed in 4% paraformaldehyde (PFA) at 4°C for 1 hour, followed by two 15-min washes with PBS at 4°C. The tumors were then chopped to 3-4 mm$^3$ chunks and incubated in anti-endomucin prepared in at least 5 volumes of PBS containing 0.5% Triton (0.5% PBT) for 6 hours at room temperature (RT) followed by incubating them at 4°C overnight. The tumor chunks were then washed in 20 volumes of 0.5% PBT for 6 hours with a change in wash buffer every hour at RT followed by a wash overnight at 4°C. The tumors were then incubated in 1:250 dilution of secondary antibodies prepared in 5 volumes of 0.5% PBT for 2 hours at RT followed by overnight at 4°C. The tumors were then washed in 20 volumes of 0.5% PBT for 6 consecutive days; for 6 hours at RT followed by overnight at 4°C each day. All steps were performed with gentle but continuous shaking. The tumors were finally cleared with 2 volumes of Vectashield (Vector; Cat: H-1000) for 2 hours at RT after which they were imaged immediately or stored in −20 °C. Steps involving antibody or Vectashield incubations were performed in 1.5-2 mL tubes and washes were performed in 50 mL tubes.

**Confocal Imaging for whole mounts**:

Tumor chunks were flattened between a 1.5 mm thick microscope coverslip (Fisherbrand; Cat: 22266858) and a double concave microscope slide (Sail brand; Cat: 7104) with their anterior walls (watershed regions) facing the coverslip. The tumors were then imaged using inverted Zeiss LSM-700 confocal microscope. Digital images of multiple z-stacks for each scanned area were captured with Zeiss Zen software and complied together using the ImageJ software. Threshold limits were set to 80% saturation and boundary of endomucin-stained channel was utilized to demarcate vasculature. Within demarcated boundary, GFP$^+$ cells were counted. Total GFP$^+$ cells were also counted. %Integrated into vasculature was determined as (GFP$^+$ cells inside vasculature*100/Total GFP$^+$ cells in field), and as an average across all z-planes per image. Stacks were analyzed individually to assess vascular integration in each plane and to reduce errors of maximum intensity projections.

**Plasmids and lentivirus:**

For knockdown experiments, the lentiviral vector used was pSicoR-GFP(Ventura et al., 2004). The sequences for the LMO2 shRNA are 5'-GACGCATTTCGGTTGAGAA-3' and 5'-GCATCCTGTGACAAGCGGATT-3'. For overexpression of LMO2 the cDNA was purchased from Genescript and cloned into the pHIV-Zsgreen vector (Addgene #18121) for lentiviral expression. Ectopic expression was verified using immunoblotting. For rescue experiments the LMO2 shRNA targeting the 3'-UTR (5'-GACGCATTTCGGTTGAGAA-3') was cloned into cloned into the pRSI12-U6-(sh)-HTS4-UbiC-TagRFP-2A-Puro shRNA expression vector (Cellecta). Viruses were produced in 293T cells using the second-generation lentiviral system and transfection using Lipofectamine 2000 (Life Technologies). Supernatants were collected at 48 and 72 hrs, filtered with a 0.45 µm filter and precipitated with Lentivirus Precipitation Solution (Alstem LLC) per manufacturer's instructions or concentrated by ultracentrifugation. Viral titers were determined by flow cytometry analyses of 293T cells infected with serial dilutions of concentrated virus.

**Xenograft tumor cell infection and engraftment:**

Dissociated single cells from xenografts were stained with biotin anti-mouse H-2Kd microbeads and depleted of mouse cells by using AUTO MACS (Miltenyi BioTec). Tumor cells were infected with pSicoR, sh*LMO2-1*, sh*LMO2-2*, at a MOI=25 for knockdown experiments for xenograft tumor. For MDA-MB-468 xenografts, cells in culture were infected at a MOI=5 and sorted for GFP expressing cells prior to xenotransplantation. MOI is calculated based on infection in 293T cells. The infected cells were washed and resuspended in staining media containing 50% Matrigel and injected in fourth abdominal fat pad by subcutaneous injection at the base of the nipple of female NSG mice (20,000 cells/mouse), except for circulating tumor cell experiment, where infected MDA-MB-468 cells were injected contralaterally in the second/third and fourth mammary fat pad. Mice were monitored every week for tumor growth. Experiment was terminated when tumors from either control or knockdown reached 1.5 cm in size or the mice showed signs of distress. Tumors were harvested, weighed, and dissociated to determine percentage of infected cells using FACS analysis. Tumor weight is plotted as 'weight × fraction infected'. The number of $GFP^+$ lung metastases were counted using the ImageJ software. In the case of PDX2, lung metastases were not discrete, so the lungs were dissociated into single cells and the percent of $GFP^+$ cells were calculated. For the tail vein injection experiment, 20,000 PDX1 tumor cells were resuspended in PBS and injected via the tail vein. The mice were closely monitored to assess any signs of distress. All the mice were euthanized 6 weeks post-injection and the lungs were harvested. $GFP^+$ lung metastases were counted using the ImageJ software.

For mouse tumors, lineage-depleted ($CD45^-$/$CD31^-$/$Ter119^-$) tumor cells from TdTomato-fluorescent *Lmo2-PyMT* were orthotopically transplanted into non-fluorescent BL6 mice. Mice were injected intraperitoneally with tamoxifen. Tamoxifen was dissolved in corn oil at a concentration of 10 mg/mL. Each mouse received a dose of 1.5 mg, as previously described (Rios et al., 2014). The mice were analyzed either 48 hrs after a single pulse or pulsed with tamoxifen 2-3 times per week once the tumors were palpable until tumor endpoint at 2 $cm^3$. At the end of the experiment, tumors were harvested, divided for histology and FACS analyzed as described above. Lungs were evaluated for metastasis and $GFP^+$ or $RFP^+$ metastases were quantified using ImageJ.

**Circulating Tumor Cells enumeration**:
Quantification of circulating tumor cells was performed in mice xenotransplanted with MDA-MB-468 cells infected with either pSicoR, sh*LMO2-1*, or sh*LMO2-2*. 400-900 μL of blood was collected via cardiac puncture using a 25-gauge needle attached to a 1-mL syringe. The blood was collected directly into K3-EDTA tubes. ACK-lysis was performed to remove the red blood cells. The cells were washed twice with FACS buffer and fixed with 2% paraformaldehyde in PBS for 10 min. The cells were again washed twice in PBS and resuspended in 100 μL of PBS. The cells were then spread on charged glass slides and allowed to air dry. Subsequently, the slides were counterstained with DAPI and the number of $GFP^+$ circulating tumor cells were manually counted under the microscope. The numbers were normalized to represent 1 mL of blood collected for each mouse.

**Migration and invasion assays:**
For migration assays, MDA-MB-468 and MDA-MB-231 cells were infected with either empty vector control or shRNA against LMO2. Subsequently, the cells were serum-starved for 24-48 hrs and 100,000 cells were plated in a trans-well dish in a 24-well plate containing 5% serum. After 36 hrs of incubation the cells in the upper chamber were removed with a cotton swab and the cells attached to the underside of the membrane were fixed in 4% paraformaldehyde. The membrane was subsequently cut and stained with 0.1% crystal violet and mounted for imaging. For 3D spheroid invasion assays, the kit was purchased from Trevigen (#3500-096-K), and the protocol was performed as per manufacture instructions.

**HUVEC Integration:**
60 μL of growth-factor-reduced Matrigel was plated per plate in a 96-well plate and allowed to gel for 30-60 mins at 37 °C. HUVEC cells were trypsinized, neutralized, and resuspended in media. 0.5 μL calcein was added per 5 mL media and incubated at 37

°C. Cells were washed once to remove calcein. 15,000 cells were plated per well onto the Matrigel coating. Once HUVEC tubes were formed at 4 hrs, 5,000 MDA-MD-468 cells transduced with either pSicoR, sh*LMO2-1*, or sh*LMO2-2* were added. Images were acquired at 8 hrs of tube formation.

**Bulk RNA-sequencing:**

200,000 MDA-MB-468 cells infected with either pSicoR, sh*LMO2-1* or sh*LMO2-2* were harvested and RNA extraction was performed using the RNeasy Micro Kit (Qiagen #74034). RNA samples were then submitted to Novogene Co. for library construction and sequencing. Briefly, following RNA quality check, mRNA was enriched using the NEBNext® Poly(A) mRNA Magnetic Isolation Module (#E7490) and cDNA libraries were constructed using the NEBNext® Ultra™ II RNA Library Prep Kit for Illumina® (#E7770, #E7775). Libraries were fragment-analyzed by LabChip, quantified by qPCR using the KAPA Library Quantification kit (#KR0405), and then sequenced on the Illumina NovaSeq 6000 platform to obtain 2 x 150 bp paired-end reads.

For data processing, raw FASTQ reads were aligned to the GENCODE v29 reference transcripts (GRCh38.p12) using Salmon (Patro et al., 2017) v0.12 with flags -l IU, --seqBias, --gcBias, --posBias, --useVBOpt, --rangeFactorizationBins 4 and --validateMapping. To calculate differentially expressed genes between control and knockdown samples, we used the R package *DESEq2 (Love et al., 2014)* (version 1.22.2) following the authors' instructions. Briefly, the gene-level count matrices were created by importing the quantification data from Salmon using *tximport* (Soneson et al., 2016) (version 1.10.1). The *DESeqDataSet* was constructed from the resulting *tximport* processed object along with sample information using the function *DESeqDataSetFromTximport*. The differentially expressed genes were calculated using the *DESeq()* function, and results summarized with the *results()* function.

Gene set enrichment analysis (GSEA) was performed on a pre-ranked list of genes differentially expressed (*Q*-value < 0.1) between control and knockdown conditions (*n*=1,963 genes) ordered by $log_2$-fold change using the Broad Institute's software (Subramanian et al., 2005). The top and bottom 50 genes from the GSEA input list were mean-centered and scaled prior to presentation as a heat map.

**Co-immunoprecipitation assay:**

Co-immunoprecipitation experiments were carried out using the Pierce Co-IP Kit (#26149, Thermo Fisher Scientific) as per manufacturer's protocol. For LMO2 and STAT3 interaction, MDA-MB-468 were grown in complete medium in a 10cm dish to reach 90% confluence. These were lysed with 0.6 mL ice cold IP Lysis Buffer containing protease and phosphatase inhibitors, for 1 hour at 4 °C upon gentle agitation. For the antibody immobilization step, 20 µg of Rabbit Anti-LMO2 (Abcam #ab91652) or 20 µg of Rabbit Anti-STAT3 CST#4904, or as a control 20 µg Rabbit IgG, were diluted onto the AminoLink Plus Coupling Resin. The cell lysates were pre-cleared with control agarose resin and co-immunoprecipitation was carried out by adding 1 mg of the pre-cleared cell lysate to the antibody immobilized resin, with end over end mixing at 4 °C overnight. After elution into 50 µL, the sample was analyzed by SDS-PAGE gel and followed by immunoblotting to detect protein-protein interaction.

**Western blotting:**

MDA-MB-468 cells were seeded, and serum starved for 24 hrs. They were subsequently treated with TNF$\alpha$ (20 ng/mL) or IL6 (20 ng/mL) for 30 mins. Whole cell lysates were generated by lysis with RIPA buffer, along with protease and phosphatase inhibitor cocktails (Thermofisher Scientific). For cell fraction the cells were lysed using the Pierce subcellular fractionation kit (PI78840). SDS-PAGE gels were run at 120V for 75 mins transferred onto PVDF membranes (#IPFL00010, Millipore, Billerica, MA) at 70V for 90min. Membranes were blocked with Li-COR blocking buffer (#927-40000) for 1 hour at RT and then subsequently probed with primary antibodies diluted at 1:1000 in 5% BSA/TBST, overnight at 4 °C. Incubation with secondary antibodies for 1 hour at RT containing fluorophores at 1:20,000 dilution (IRDye 800CW conjugated goat anti-rabbit #926-32211, LI-COR Biosciences, Lincoln, NE) enabled visualization on the Odyssey Infrared Imaging System from LI-COR Biosciences. Washes in between incubations were done for 10 mins x 3 using TBS-Tween 0.1%.

**Luciferase reporter assay:**

STAT3 firefly luciferase reporter lentivirus (PLV-10065-50) was purchased from Cellomics technologies. A stable cell line with MDA-MB-468 was generated using puromycin selection. The cells were then infected with either pSicoR, sh*LMO2-1* or sh*LMO2-2*. For the reporter assay, 10,000 MDA-MB-468 cells were seeded in full serum media. After attachment, there were treated with TNF$\alpha$ (20 ng/mL), IL6 (20 ng/mL) or EGF (10 ng/mL) in full serum for 4 hrs. The cells were then lysed and luciferase activity was measured Dual-Luciferase Reporter Assay System (Promega, #E1960). All fold changes were calculated based on untreated cells in the same group i.e pSicoR, sh*LMO2-1* or sh*LMO2-2* that are untreated. All experiments were performed in triplicate and the experiment was repeated 3 times.

**DUOLink Proximity Ligation Amplification (PLA) assay:**

For the PLA assay (DUOLink, OLink Biosciences, Sigma-Aldrich #DUO92102), MDA-MB-468 were seeded on 13 mm glass coverslips. The cells were fixed with ice cold 100% methanol for 5 mins at −20 °C and then rehydrated thrice in PBS for 5 mins each. Coverslips were blocked for 30 mins with 3% BSA/PBS and then incubated with appropriate dilution of primary antibodies in 1% BSA/PBS for 1 hour in a moist environment at room temperature. Primary antibodies were used at 1:200 dilutions to characterize the interaction between LMO2, STAT3, JAK2 and PIAS3. All antibodies are listed in Table S5. As a negative control, rabbit and mouse anti-IgGs were used in 1:200 dilutions. Subsequently, manufacturer's instructions were followed to complete the PLA assay.

**Statistical analysis:**
All graphs show the average as central values and error bars indicate ± SD unless otherwise indicated. *P*-values are calculated using paired or unpaired *t*-test, ANOVA, ELDA, Wilcoxon Rank-sum test, Monte Carlo simulation, Fisher's exact test, and log-rank test as indicated in the figure legends. All *P*- and *Q*-values were calculated using Graphpad prism or R version ≥ 3.5.2, unless otherwise stated. For animal studies, sample size was not predetermined to ensure adequate power to detect a pre-specified effect size, no animals were excluded from analyses, experiments were not randomized, and investigators were not blinded to group allocation during experiments.

**Data code and availability statement:**
All data generated or analyzed during this study are included in this published article (and its supplementary information files). Source data for figure files are available and will be provided on request. All single-cell and bulk RNA-sequencing data generated in this study have been deposited in the Gene Expression Omnibus with the primary accession code GSE159285 (https://www.ncbi.nlm.nih.gov/geo/query/acc.cgi?acc=GSE159285). All bioinformatics tools used in this study are published and publicly available. Custom scripts are available from the authors upon reasonable request. Patient-derived xenografts generated in the manuscript will be available upon request with an appropriate Material Transfer Agreement (MTA) with Stanford University. Requests for the *Lmo2*$^{CreERT2}$ mice should be addressed to Dr. Terence Rabbitts.
